# Systematic assessment of regulatory effects of human disease variants in pluripotent cells

**DOI:** 10.1101/784967

**Authors:** Marc Jan Bonder, Craig Smail, Michael J. Gloudemans, Laure Frésard, David Jakubosky, Matteo D’Antonio, Xin Li, Nicole M. Ferraro, Ivan Carcamo-Orive, Bogdan Mirauta, Daniel D. Seaton, Na Cai, Danilo Horta, HipSci Consortium, iPSCORE Consortium, GENESiPS Consortium, PhLiPS Consortium, Erin N. Smith, Kelly A. Frazer, Stephen B. Montgomery, Oliver Stegle

## Abstract

Identifying regulatory genetic effects in pluripotent cells provides important insights into disease variants with potentially transient or developmental origins. Combining existing and newly-generated data, we characterized 1,367 iPSC lines from 948 unique donors, collectively analyzed within the “Integrated iPSC QTL” (i2QTL) Consortium. The sample size of our study allowed us to derive the most comprehensive map of quantitative trait loci (QTL) in pluripotent human cells to date. We mapped the effects of nearby common genetic variants on five expression phenotypes, identifying *cis*-QTL at gene-, exon-level and transcript-, splicing-, alternative polyadenylation-ratio (APA) for a total of 18,556 genes. For gene-level, we further quantified the effects of rare and singleton variants, and the effect of distal variants that act in *trans* (*trans*-eQTL), which we replicated in independent samples. Our data are a valuable community resource, uncovering novel regulatory effects that have not previously been described in differentiated cells and tissues. Building on this regulatory map, we functionally explore GWAS signals for over 4,336 trait loci, finding evidence for colocalization with common and rare iPSC QTL for traits such as height and BMI, and diseases, such as cancer and coronary artery disease.

## Introduction

Genome-wide association studies (GWAS) have yielded a compendium of genetic variants that are associated with human traits and diseases. However, the majority of these variants are in intergenic regions^1^, and identifying their molecular function remains an ongoing challenge. Molecular quantitative trait loci (QTL) studies have been performed to systematically identify the function of genetic variants via their effects on molecular intermediates such as gene expression levels. However, existing QTL resources are largely focused on differentiated cell types, including blood^2–4^ and post-mortem collected tissues^5^. In contrast, much less is known about the impact of genetic variants on molecular intermediates in pluripotent cells, which have only recently become accessible via the first population-based iPSC studies with moderate sample sizes^6–10^. Due to their embryonic-like state, iPSCs can provide unique insights into genetic regulation of expression in cell states that mimic early development, with relevance to diseases that manifest *in utero* or in transient states throughout development^9,11,12^.

We characterized 1,367 iPSC lines from 948 unique donors, collectively analyzed within the “Integrated iPSC QTL” (i2QTL) Consortium, and provide a comprehensive map of expression regulatory variants in human iPSC. We mapped the effects of nearby common genetic variants on five expression phenotypes, identifying *cis*-QTL at gene-, exon-level and transcript-, splicing-, alternative polyadenylation-ratio (APA). For gene-level, we further quantified the effects of rare and singleton variants, and the effect of distal variants that act in *trans* (*trans*-eQTL). Our data are a valuable community resource, uncovering novel regulatory effects that have not previously been described in differentiated cells and tissues. Building on this regulatory map, we functionally explore GWAS signals for over 4,336 trait loci, finding evidence for colocalization with iPSC QTL for traits such as height and BMI, and diseases, such as cancer and coronary artery disease.

## Results

Within the i2QTL Consortium we collected previously published and novel data from human iPSC lines (**Table S1 & S2**), providing genotype and RNA-seq data from a total of 1,367 iPSC lines, derived from 948 unique donors. The data were obtained from five iPSC consortia; iPSCORE^6^ (N=206), GENESiPS^10^, (N=85), PhLiPS^7^ (N=65), Banovich *et al.*^8^ (N=49) and HipSci^9^, which has been greatly expanded in size for this study (N=543, original study N=166). Additionally, data from the Choi et al^13^ study were included as reference, adding a total of 92 samples comprising fibroblast, embryonic stem cell (ESC), and iPSC (**Table S1 & S2**).

In order to mitigate technical differences, we uniformly reprocessed all genotype data, considering both genotyping array and whole genome sequencing data, as well as uniformly reprocessed all RNA-seq data (see Methods). Joint multidimensional scaling (MDS) of i2QTL and GTEx^5^ (v7) samples revealed a remarkable level of homogeneity between the iPSC studies compared to between-sample and tissue differences observed in GTEx, with the remaining variability in iPSCs appearing to being primarily due to differences in sequencing protocol between iPSC consortia (**Fig. 1a**). Using samples from Choi et al^13^, we were able to confirm both the minimal processing effects, using the fibroblast lines, and, using ESC lines, the embryonic-like state of our iPSC lines (**Fig. 1a**).

**Figure 1.**
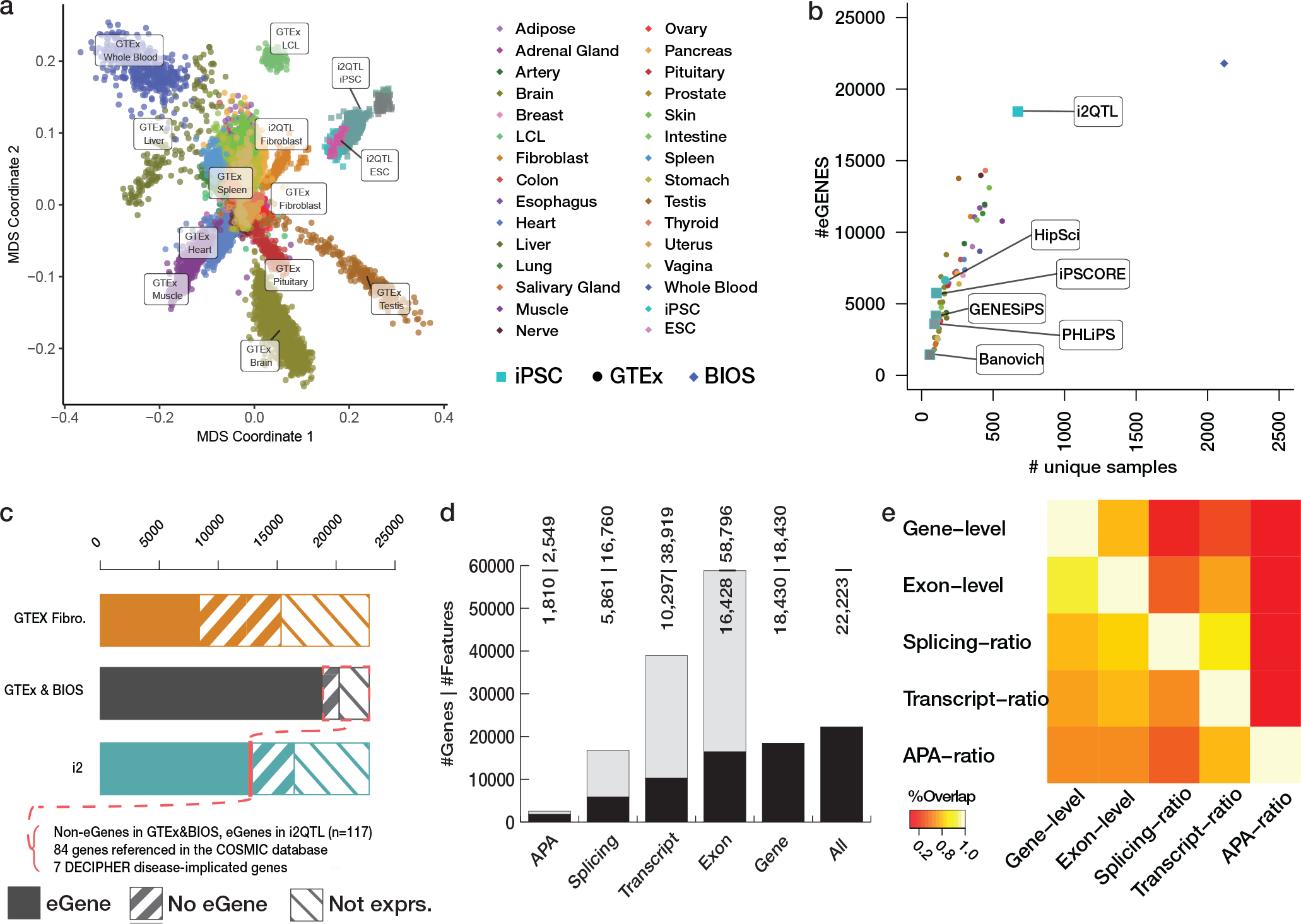
Discovery of *cis* genetic regulation across a comprehensive set of RNA phenotypes. **A**. Comparison of gene expression profiles of iPSC vs the GTEx (v7) tissues. Shown are the first two MDS components obtained from gene expression levels. **B**. Comparison of the number of discovered eGenes as a function of sample size for existing iPSC studies, GTEx (v7) and BIOS. **C**. Comparison of eQTL coverage for protein-coding genes in Fibroblasts (orange), the combination of the GTEx tissues (all results shown in GTEx v7), BIOS blood eQTL (black) and i2QTL iPSC eQTL (blue). The fraction of protein coding eGenes without previous evidence for eQTL in BIOS & GTEx is shown in red (see Results). Disease association is shown at the bottom. **D**. Breakdown of the identified *cis*-QTL per expression phenotype. Bar plot, displaying both the number of associations for individual phenotypes (grey) and the number of genes with at least one association (black) per tested feature per expression phenotype (eFeature) and associated gene (#eGenes | #eFeature). **E**. Pairwise replication of genetic effects between phenotypes, showing the fraction of replicated QTL loci. Shown are the fraction *cis*-QTL discovered at FDR 5% discovery (rows), and replicated at FDR 10% (columns).

### Mapping *cis* regulatory effects in iPSCs

We mapped *cis*-QTL, considering proximal (within the gene body +/−250kb) common variants (MAF >1%) and paired-end stranded RNA-seq data available for 936 samples (N=682 donors) of European ancestry (see Methods).

Among 27,046 tested Ensembl genes, we identified at least one *cis*-eQTL for 18,430 genes (68% of tested; FDR<5% permutation-based, **Table S3**, Methods; in the following denoted eGenes), which corresponds to a 2.5 fold increase in the number of *e*Genes compared to the largest previous *cis*-eQTL map in iPSCs^9^ (**Fig. 1b**). Conversely, the vast majority of the eQTL effects (between 62-86%) described in the previous studies were detected in our data (replication defined at nominal P<0.05 with matched effect direction) **(Fig. S1-4)**. We also performed iterative eQTL mapping using stepwise regression (see Methods), which identified two or more independent effects for 39.0% of eGenes, with up to 12 independent *cis*-eQTL for *PTGR1* **(Fig. S5)**.

To gain insights into iPSC-specific regulation, we contrasted our *cis*-eQTL map with somatic eQTL maps from GTEx^5^ tissues and blood eQTL from BIOS^4^. We observed a greater number of eGenes in iPSC as compared to any of the single GTEx tissues (**Fig. 1b-c**), and approximately equal numbers as observed in BIOS, despite the substantially larger sample size of this study (**Fig 1b)**. The increase in power to detect eGenes in iPSC is in line with previous results^9,14^, likely reflecting the homogeneity of the iPSC lines^15^. We identified 994 eGenes without prior evidence for regulatory effects in neither GTEx nor BIOS (**methods**; **Table S3**), 117 of which are protein-coding (**Fig 1c**). This set of iPSC-specific eGenes are enriched for known cancer genes curated in the COSMIC database (p<2.2×10^−16^, 84 novel eGenes are in COSMIC), and we identified eQTL for seven genes in the DECIPHER rare disease database (**Fig 1c**).

We extended the widely considered gene-level eQTL mapping to an extended set of expression phenotypes (transcript-ratio, exon-level, splicing-ratio (sQTL) and APA-ratio) (**Table S4-7**). To aid comparability, we considered the same gene-based region to test a consistent set of variants, followed by gene-based FDR adjustment (**methods**). In aggregate, across all five expression phenotypes, this identified genetic effects for 18,556 Ensembl genes (FDR<5% permutation-based, Methods; **Fig 1d**), where 77% of the eGenes discovered in one phenotype were also detected when assessing at least one other expression phenotypes. Despite this overlap, we observed 3,118 genes (9.8% of all tested genes) with an exon-, transcript-, splicing- or APA-QTL but no gene-level eQTL, highlighting the benefits of our multi phenotype QTL mapping strategy. Assessment of the pairwise replication between these maps at the level of individual QTL variants confirmed substantial sharing between phenotypes, with the largest overlap identified between exon-level and gene-level QTL, and most distinct for APA-ratio QTL (**Fig 1E**).

### Mapping rare variant effects on gene expression

Rare coding and non-coding variants are associated with comparatively larger effects on gene expression than common variants^16,17^, which contributes to increased disease risk^18,19^. As the frequency of variants that can be assessed using conventional QTL mapping is limited by sample size, we sought to identify the landscape of rare genetic variants linked to outlying expression profiles of individual genes within iPSCs.

We considered the subset of lines with both RNA-seq and WGS available (N = 425 cell lines after filtering - Methods, **Table S2**), and focused on variants within, or up to 10 kb around, protein-coding and long non-coding RNA genes. Gene expression outliers were defined as the sample with minimum gene expression Z-score (Z-score < −2, under-expression outlier) or maximum gene expression Z-score (Z-score > 2, over-expression outlier) for a given gene. We computed variant burden scores, equivalent to relative risk (**methods**), across genes and assessed differences in variant burden in outliers compared to non-outliers, aggregating across gnomAD MAF thresholds (both common and rare) and deleterious annotation (CADD - where available). This score was computed separately for gene-level under-expression outliers and over-expression outliers. Notably, for both SNPs and indels, we observed an enrichment in under-expression outliers for rare (gnomAD MAF 0<MAF<=0.01%), highly-deleterious (CADD>25) variants (5-fold and 40-fold increase in comparison to non-outlier in SNP and indels respectively) (**Fig 2A**) (**Table S8**). For structural variants (SVs), a 9-fold increase in rare (study MAF 0<=MAF<1%) duplications, and 18-fold increase in rare multi-allelic copy number variants (mCNVs), was observed in over-expression outliers compared with non-outliers (**Fig 2B**) (**Table S8**). The findings for SVs is in line with recent findings detailed in Jakubosky et al^20^.

**Figure 2.**
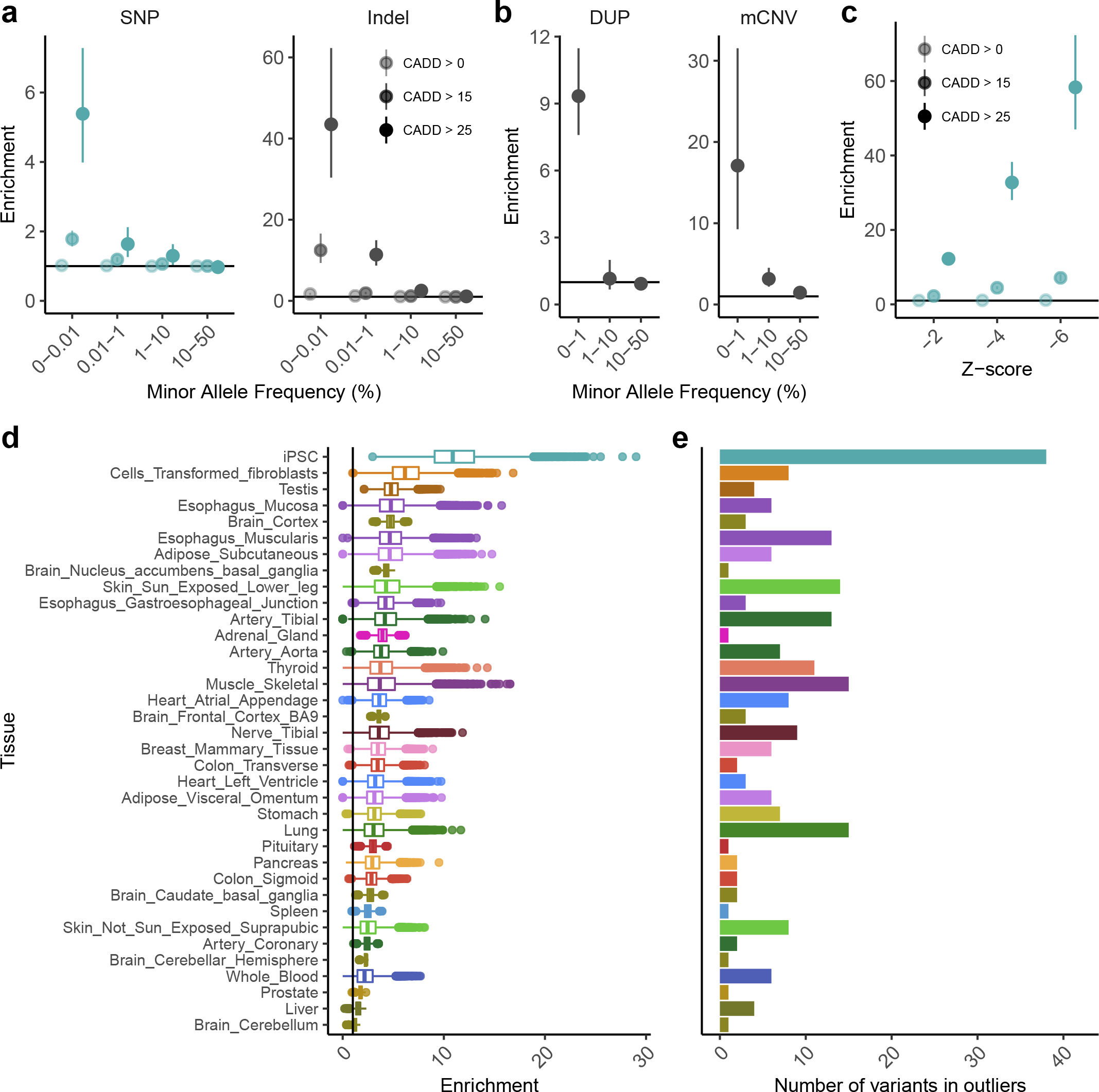
Linking rare variants to gene expression outliers. **A.** Enrichment of deleterious rare SNPs and indels in samples with gene expression outliers; **B.** Enrichment for rare duplications and mCNVs in gene expression outliers. **C.** Enrichment for singleton, deleterious SNPs in gene expression outliers, identifying stronger enrichments in under-expression outliers. **D.** Comparison of enrichments for singleton, highly-deleterious (CADD > 25) SNPs in iPSC (panel C, Z-score = - 2, dark-green point) versus GTEx v7 tissues (adjusted for differences in sample size; Methods). Shown are enrichment scores for 10,000 random draws of 50 samples. Enrichment is strongest in iPSC. **E.** Highest proportion of outlier effects for shared set of rare, deleterious variants (i.e. variants passing thresholds and found in both i2QTL iPSC and GTEx) are in iPSC.

To further identify the extent to which the frequency of a variant in the studied population is associated with outliers compared to non-outliers, we computed enrichment scores for singleton SNPs (where a singleton is defined as a SNP not observed in gnomAD (**Fig 2C**)). This approach allows for assessing the full frequency spectrum of SNPs, from ultra-rare to common. The analysis was repeated for increasing outlier stringency, from Z-score −2 to −6. Singleton, high-CADD SNPs (CADD>25) were found at a 12-fold higher rate in under-expression outliers than non-outliers at a Z-score of Z <− 2 (up to 60-fold when Z < −6). The estimate for singleton, high-CADD (CADD>25) SNPs (12-fold increase) is substantially higher than that observed for the next variant bin at the same CADD and Z-score threshold (5-fold increase) (**Fig 2A**).

To place these enrichments into context with somatic tissues, we repeated the outlier analysis for singleton, high-CADD (CADD>25) SNPs in 35 tissues from GTEx (v7) with 50 or more samples remaining after global outlier removal on PEER adjusted data^21^ (**Fig 2D**; Methods). To control for variation in sample size across tissues, we calculated the enrichment across 10,000 random draws without replacement of equal sample size for each tissue (N samples = 50). In this comparison, iPSC had the greatest enrichment score (median ~9), followed by GTEx fibroblasts (enrichment ~7) and GTEx testis (enrichment ~5) (**Table S9**). The observed enrichment was moderately correlated with number of genes expressed in each tissue (Pearson correlation = 0.34); however, when randomly subset to a fixed number of genes across tissues, the maximum median enrichment was still observed for iPSC compared to GTEx tissues (**Sup Fig 6**).

Motivated by the hypothesis that rare variants are driving observed gene outlier effects, we considered the variant class with the largest burden scores (i.e. gnomAD singleton, CADD>25 SNPs) to study the intersection of these variants present in both GTEx and i2QTL samples (N variants = 305), thereby assessing the relative discovery of outlier effects in genes linked with a shared set of ultra-rare, deleterious variants. Specifically, this analysis helps to clarify the power of rare variant-driven outlier effects, when explicitly controlling for potential differences in variant calling between i2QTL and GTEx. In total, 38/305 (12.5%) of shared SNPs were identified in outlier samples in iPSC, with the next closest being GTEx skeletal muscle and lung outlier samples (N variants = 15) (**Table S10**). Furthermore, 20 of the 38 SNPs observed in iPSC outlier genes were not observed in outliers across any of the 35 GTEx tissues selected (i.e. the variants were observed in outlier samples in iPSC only, and in non-outlier samples across all GTEx tissues). In GTEx, these variants were associated with an overall median absolute Z-score across tissues of 0.49 (range 0.27-1). Four of these i2QTL-only outlier genes have Developmental Process annotations in Gene Ontology (GO), and six have known OMIM disease associations (**Table 1**).

**Table 1.**
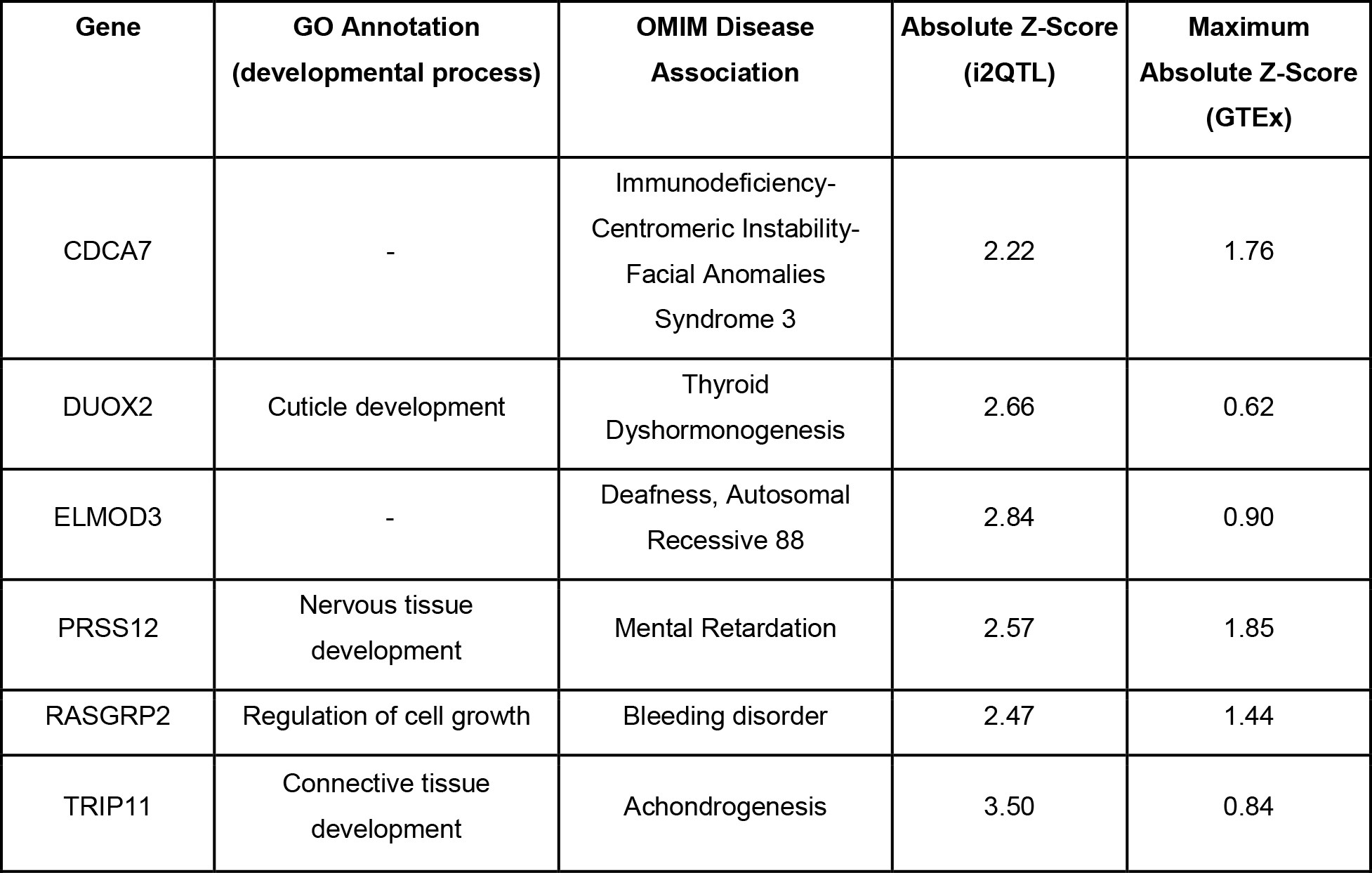
Genes linked to singleton, highly-deleterious SNPs with transcriptome outlier effects in pluripotent cells only

### Distal genetic effects linked to gene expression

Next, we leveraged the full dataset of individuals from European ancestry (samples N=1,123, donors N=759, Table S2 see Methods) to probe for downstream regulatory effects that act in *trans* (defined as >2.5 mb away from the gene body). The identification of *trans*-eQTL is challenging due to the large number of tests and the typically smaller effect sizes, and hence only feasible in large cohorts. To mitigate the multiple testing burden we considered *trans* effects on protein-coding genes (average TPM>1; 16,451 genes), assessing *cis*-QTL variants (N=93,146), originating from the *cis*-QTL maps described above, as well as known GWAS variants (N=23,798; NHGRI-EBI GWAS catalog^22^ v92), resulting in a total of 115,709 unique variants in our analysis.

Genome-wide, this identified 193 independent *trans*-eQTL comprising 191 unique genes (FDR<10%; permutation-based adjustment, **Table S11 & S12**, Methods). The majority of *trans*- eQTL were exclusively linked to *cis*-QTL variants (163, 85.5%), with 21 *trans*-eQTL linked to both GWAS and *cis*-QTL variants and seven *trans*-eQTL exclusively linked to a GWAS variants. Notably, 138 of the 184 *cis*-QTL with *trans* effects had *cis* effects originating from more than one gene phenotype in *cis*. The largest number of effects downstream of non-gene-level *cis*-QTL were observed for transcript-ratio *cis*-QTL (33 of the 46). Notably, 46 *trans*-eGenes (24% of the *trans*-eGenes) would not have been identified when restricting the analysis to conventional gene-level *cis*-eQTL and/or GWAS variants. Collectively, these results highlight the relevance and the potential for downstream consequences of *cis*-QTL that do not act on a gene level.

To assess the functional relevance of the identified *trans*-eQTL, we focused on four *trans*-eQTL hotspots, defined as *trans*-eVariants, or blocks of *trans*-eVariants within 250Kb, associated with five or more unique genes. The hotspot with the largest number of downstream genes affected was located near *ELF2*, a transcription factor linked to reduced proliferation^23^. The hotspot comprises effects of 16 individual *cis*-QTL variants for *ELF2* and genes in the vicinity; these *cis*-QTL effects could be decomposed into three major blocks (between block LD < 0.67) (**Fig. 3C**). Variants in the different blocks were linked to multiple *cis*-QTL types, mostly on *ELF2*, but also for APA ratio change of *NAA15* (**Fig. 3D**). Despite moderate LD between variants, we observed a high degree of sharing of downstream *trans*-eGenes between *cis*-eVariants (**Fig. 3E**). The 37 downstream genes, linked to the ELF2 hotspot, were enriched for sequence motifs from the ELF transcription factors, which have a strong sequence resemblance. Significant associations were observed for *ELF3*, *ELF4* and *ELF5* (p-adj: 3.3×10^−7^, 6.8×10^−5^ and 2.2×10^−6^ respectively, g:Profiler^24^), and 20 out of the 37 *trans*-eGenes were also known target of *ELF2*. A second hotspot, in *cis* linked to a *CREB3L2* with *trans* effects on 8 eGenes, did stand out because of an enrichment for the reactome pathway: “ER to Golgi Anterograde Transport” (p-adj<5.6×10^−5^), which is consistent with previous associations between *CREB3L2* and both the ER and Golgi complex^25^.

**Figure 3.**
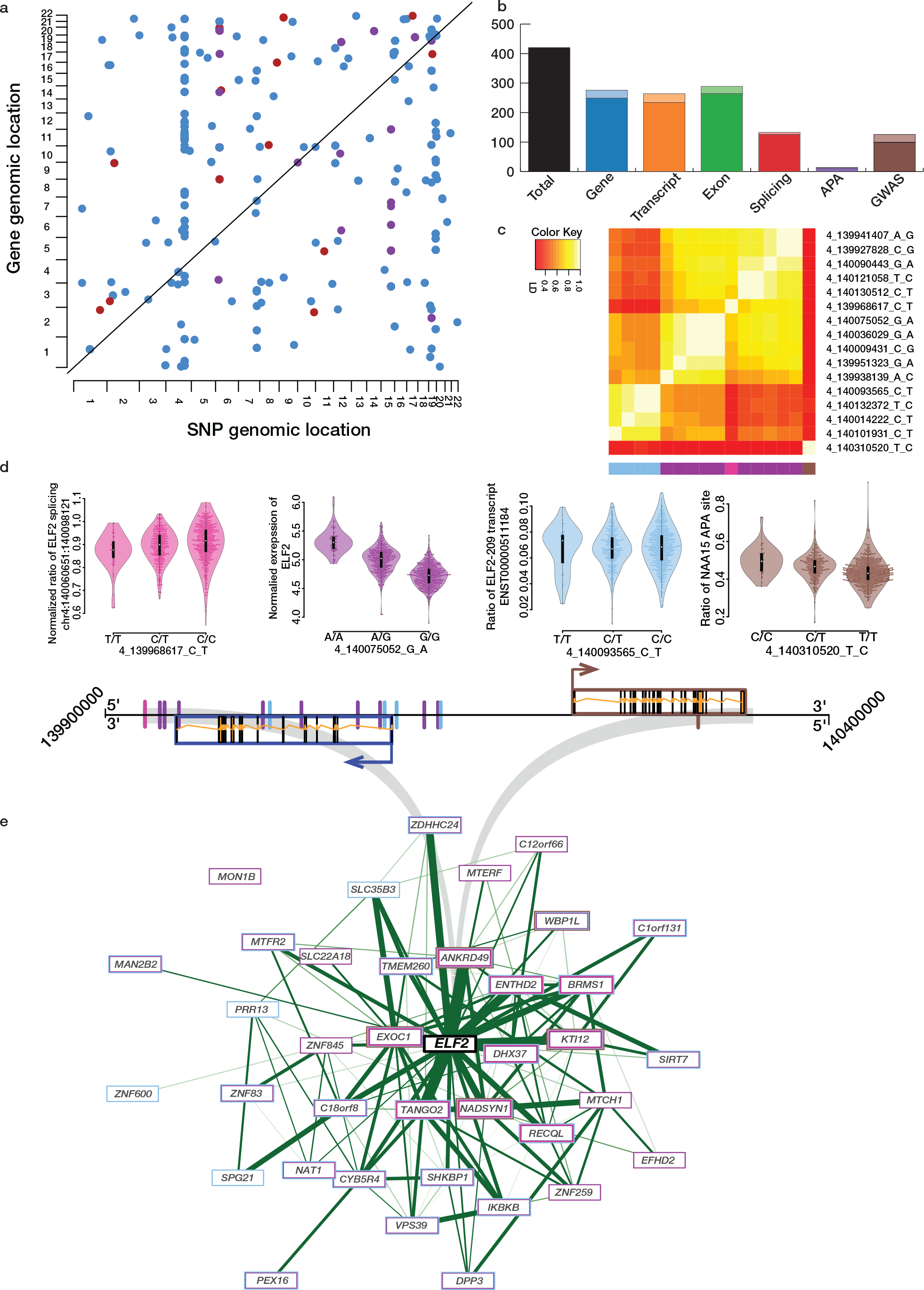
Discovery of distal (*trans*) genetic regulation on gene expression. **A**. Dot plot of the independent *trans*-eQTL in iPSC (FDR<10%). Each dot corresponds to one *trans*-eQTL, with color denoting variant category (blue: *cis*-QTL, red: GWAS variant, purple: shared effects). **B**. Total number of *trans*-eQTL per variant category, displaying the total number of effects. Darker shaded colors denote *trans*-eQTL linked to multiple variant types, the lighter shades correspond to the number of unique effects for a given variant category. **C-E** Dissection of the *trans*-eQTL hotspot near *ELF2*. **C.** LD structure between *cis* eQTL variants implicated in the hotspot, indicating three major and one minor effect groups linked to specific *cis* eQTL (blue: transcript-ratio effect on *ELF2*, purple: *cis*-eQTL on *EFL2*, pink: splice effect in *ELF2,* brown: the APA effect on *NAA15*). cis-QTL effects of the four variants identified in **D**, acting on different gene phenotypes of *ELF2* (left) and *NAA15* (right). SNPs linked to effects in the regions are shown on the illustration colored based on the effect they tag. **E**. The co-expression network of the genes that are controlled by the *trans*-eQTL linked to the hotspot near *ELF2*, including *ELF2* (in the center). The genes are color coded by the *cis* QTL variant that drives the *trans* effect (colors as in **C** and **D**). Genes for which multiple regulators were identified are depicted using multiple colored rings around the gene name, color of the ring based on QTL effects in **D**.

We identified 28 *trans*-eQTL variants linked to GWAS variants, including variants linked to breast cancer, congenital craniofacial abnormalities, age of menopause and metabolic traits (Supplementary note 1). For example, the most significant *trans*-eQTL (p-adj=1.08E-15) associated changes in expression of NBPF14 to a GWAS hit (rs11249433) on breast cancer; NBPF14 was previously identified as one of the most frequently mutated genes in breast cancer^26^. The same GWAS variant was also a *cis*-QTL for three (breast) cancer-implicated genes FAM72^27^, RP11–439A17.7^28^, HIST2H2BA^29^. Another interesting example was found for cleft palate, cleft lip and hemifacial microsomia, with a set of four variants (rs11072494, rs2289187, rs10459648, rs6495117) linked to six downstream *trans*-eGenes (*GPR160*, *SEMA3A*, *MDFIC*, *GRIK2*, *FAM169A* and *GLB1L3*). This set of genes is enriched for the JNK cascade and the MAPK cascade biological processes (g:Profiler p-adj: 1.0×10^−2^, 2.1×10^−2^ respectively), with known implications in congenital craniofacial abnormalities^30,31^.

To assess the validity of our set of *trans*-eQTL, we used the remaining iPSCs from non-European samples (N=187), and assessed their marginal replication rate in these independent samples. In total, 17% of the effects could be replicated (nominal P<0.05 and matched effect direction; Methods); a replication rate that exceeds chance expectation (median replication rate 5.3% assessed by testing random *trans*-eQTL pairs (see Methods)). We also sought to replicate our trans association in other cell types using existing eQTL resources, however none of the effects replicated in blood^2^ or any of the GTEx^5^ tissues. However, 12 *trans*-eQTL SNPs that a in strong LD (LD>0.8) with previously published *trans*-eVariants, although affecting different genes (**Table S12**). This result is consistent with reports from GTEx, showing substantial heterogeneity in *trans* eQTL across tissues^5^. To further assess the robustness of the *trans*-eQTL, we tested the cross-omic replication of *trans*-eQTL in DNA-methylation data, available for a subset of the samples (N=572 donors, N=841 lines), finding evidence for cross-omic replication for 27% of the identified *trans*-eQTL (**methods**), again exceeding the chance expectation (median replication rate of *trans*-meQTL: 15.3; **methods**). When combined, we find replication evidence for 38% of the identified *trans*-eQTL, replication of the hotspot *trans*-eQTLs was higher (45.3%, Fisher exact P-value 0.01; **Table S12**).

### Colocalization of GWAS signals and molecular QTL

A major opportunity provided by eQTL maps in pluripotent cells is the identification of disease-associated variants that colocalize with eQTL, providing evidence for a developmental or transient mechanism. Using a combination of FINEMAP^32^ and eCAVIAR^33^ (see Method), we systematically assessed colocalization between the presented *cis*-QTL map, comprising of QTL for five gene phenotypes, and a broad range of previously reported genetic associations obtained from diverse GWAS studies (N=350), collected from the Phenome Scanner Database V2^34^, the NHGRI-EBI GWAS catalog^22^, GWAS curated in LocusCompare^35^, and 1,740 traits from the UKBB phase 1 GWAS^36^.

In total, we identified 4,336 colocalization events (**methods**), linking 608 disease- and phenotype loci to 10,794 *cis*-QTL loci, which span the full range of expression phenotypes (**Table S13**). These events covered a broad range of traits and disease, with the largest number of individual colocalizations linked to physical traits, including height and BMI, lab-measurements, including blood-cell (composition) traits, and diseases such as Crohn’s disease and coronary artery disease (CAD) (**Fig 4A**). Notably, when restricting this analysis to gene-level eQTL, over 40% of the colocalization events are lost. The largest number of QTL-type specific colocalization events were observed for the exon-level QTL, followed by colocalization events with gene-level and transcript-ratio QTL (**Fig 4B**). As an illustration for CAD, 36 out of 93 GWAS loci were found to colocalize with an iPSC QTL, with at least one QTL-type specific colocalization event that would not be detected by any of the remaining QTL types (**Sup Fig S7**). Next to the *cis*-QTL colocalization we also test for co-localization between *trans*-eQTL and GWAS loci, and finding, amongst others, significant overlaps betweenthe *ELF2* and *CREB3L2* hotspots with height (**methods**; **Supplementary note 1**; **Table S14**).

**Figure 4.**
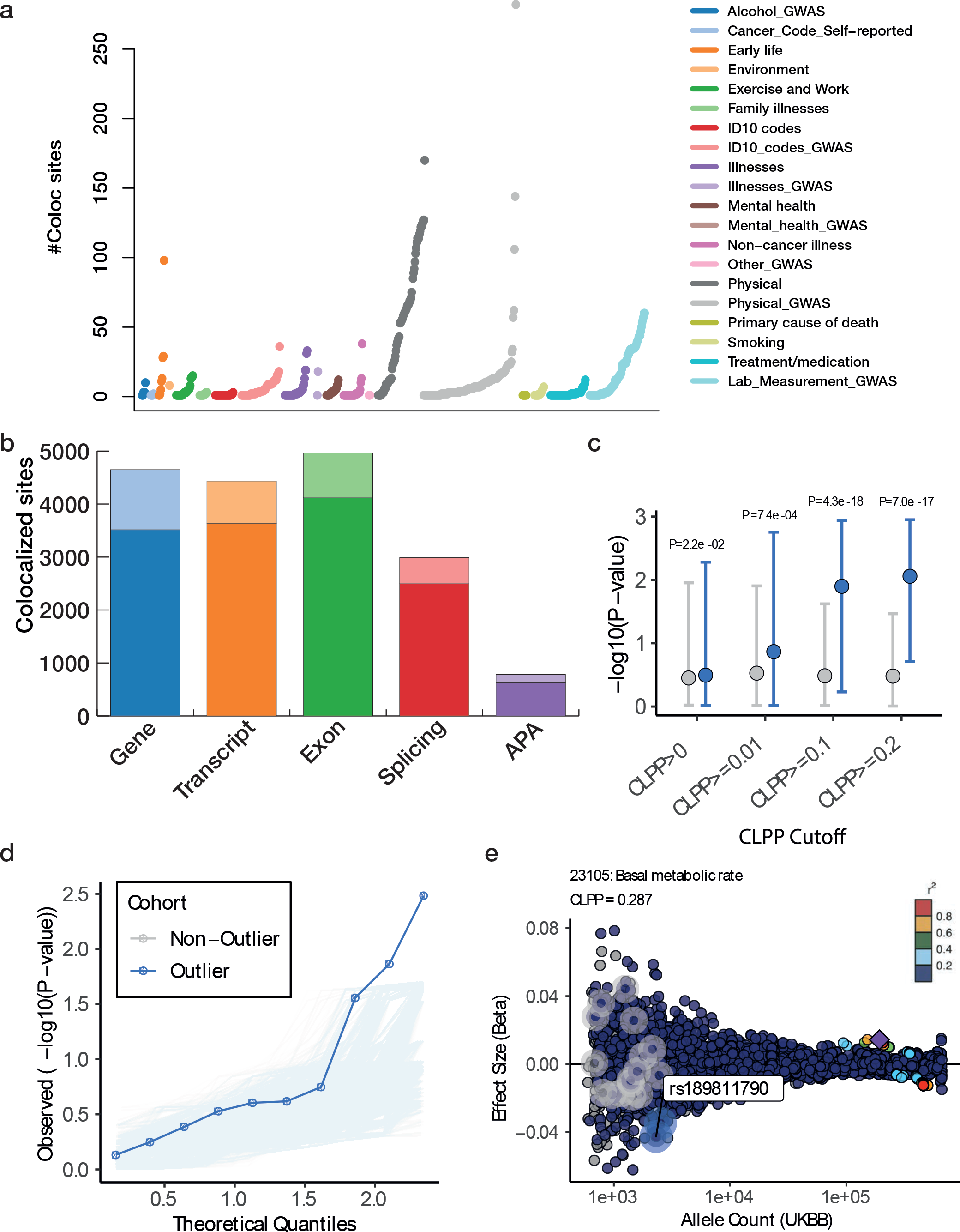
Colocalization of disease and traits variants with iPSC QTL, and associations with expression outliers. **A**. Overview of the number of colocalization events for individual GWAS, grouped by trait categories (based on UKBB). **B**. Summary of colocalization events for individual *cis*-QTL type, displaying the total number of GWAS loci with colocalization events that are specific to a QTL type (light colour; not detected by any other QTL type) or shared with at least one other QTL type (dark colour). **C**. Negative log P-values of GWAS trait associations (mean, +/− 95% CI) for iPSC outlier- versus non-outlier-associated variants, considering genes with varying degree of evidence for co-localization with common eQTL variants. Genes are stratified by the colocalization posterior probability (CLPP) score. Outlier-associated variants had overall greater effects in GWAS in which there was evidence for co-localization of the same genes, compared to matched non-outlier variants, increasing with colocalization probability. **D**. Negative log P-value of for variant associations with basal metabolic rate (UKBB GWAS ID: 23105), considering outlier- (blue line) and non-outlier-associated (gray lines) variants. Non-outlier-associated variants were chosen randomly to match the number of outlier-associated variants (repeated 10,000 times). The associations of outlier-associated variants were more significant than the observed associations of any non-outlier combination sampled. **E**. Example of a gene locus showing GWAS effect size for outlier-associated variants (blue highlight) and non-outlier-associated variants (gray highlight). The greatest protective effect sizes are observed for outlier-associated variants. Points are colored by LD (1000 Genomes European samples) relative to lead variant (top P-value) (purple diamond) in gene locus.

As some GWAS, such as BMI or height, have a large number of GWAS hits reported, we examined the fraction of GWAS loci that colocalize with QTL, considering traits with at least ten loci that were tested for colocalization. The highest relative overlap was found for primary biliary cirrhosis (PBC), followed by triglyceride (TG) levels. For PBC, 10 out of the 12 reported GWAS loci colocalize with *cis*-QTL effects. The corresponding genes were enriched for MAPK, NF-kappa B and TNF-R1 signaling pathways (g:Profiler p-adj: 1.1×10^−2^, 4.9×10^−2^, 5.725×10^−3^ respectively), with known functions in the immune system. For TG levels, 12 out of 15 reported GWAS loci overlap with iPSC QTL. Enrichment analysis for these QTL showed significant overlap with metabolic pathways (g:Profiler: alpha-Linolenic acid metabolism, p-adj: 4.899×10^−3^; Biosynthesis of unsaturated fatty acids, p-adj: 5.729×10^−3^, Fatty acid metabolism, p-adj: 2.585×10^−2^). Furthermore, even when normalizing by the total number of loci linked to a GWAS trait, many physical traits still appeared as top colocalized traits.

Integration of transcriptomic data with GWAS summary statistics offered the opportunity to understand the distribution of large-effect rare variants across many diseases and traits. Given the sample size of large scale GWAS, such as UKBB, rare variants become testable in GWAS; however, due to the (at present) large discordance in sample size when assessing genetic effects on expression we cannot apply standard eQTL mapping to finemap causal rare variants. Here we integrated information on gene colocalization and gene outlier expression levels to annotate function to rare variants. To assess the impact of rare variants on GWAS traits we considered variants linked to iPSC outlier- and non-outlier samples, for which summary statistics were available in UKBB Phase 1 GWAS. We first selected variants observed within one sample in i2QTL only; specifically, each variant was tabulated across all samples, and variants appearing more than once were removed from the analysis. For outlier-associated variants, this has the effect of isolating the set of variants putatively driving observed outlier gene expression (i.e. should the same variant be observed in both an outlier and non-outlier sample, by definition this would suggest that the variant is less likely to be causing the observed outlier expression). In addition, we filtered for variants with a MAF of 0<MAF<=1% from gnomAD and considered genes for which we observed evidence for co-localization with the GWAS trait with common variants (CLPP score > 0). For each outlier variant, we also considered a matched non-outlier variant linked to the same gene (matched CADD score in a range +/− 5, gnomAD MAF 0<MAF<=1%). Given that UKBB Phase 1 GWAS used imputed genotypes from common variants measured via microarray for rare variant calling^37^, the intersection of variants was considerably smaller than the set of donor-specific rare variants in i2QTL. After intersecting these datasets, we obtained 10,103 outlier- and non-outlier associated variants linked to 779 genes and 2,419 traits.

From this set of variants, we investigated genes with at least one outlier and one non-outlier variant in/near the gene (N traits = 543) and compared the GWAS p-values from outlier- and non-outlier-associated variants with increasing evidence of trait colocalization CLPP cutoffs (**Fig 4C**). Genes that colocalized with the QTL results suggest that variation in the gene locus is involved in modulating risk for a given trait. For increasing CLPP score from CLPP>0 to CLPP>=0.2, we observed overall more significant GWAS p-values for outlier-associated variants than non-outlier-associated variants (CLPP>0 p=0.02; CLPP>=0.2 p<1e−16), a trend that was also observed for the effect size estimates (data not shown). To examine individual traits, we also considered a quantile-quantile plot to compare the empirical distribution of outlier-associated variant p-values in GWAS compared to matched non-outlier variants. For example, for basal metabolic rate (UKBB GWAS ID: 23105), this yielded an inflation in outlier variant p-values compared to 10,000 random, matched selections of non-outlier variants (**Fig 4D**). Across all outlier-associated and randomly-selected non-outlier variants (shown in **Fig4D**), the top variant p-value was observed in the gene *HSD17B12*, a gene known to be involved in type 2 diabetes mellitus^38^, and with a CLPP score of 0.29 in the present study (**Fig 4E**). Notably, this specific outlier variant has one of the largest GWAS effect sizes, when considering a 1Mb locus, but owing to its frequency this variant does not pass statistical significance threshold in conventional GWAS analysis. Overall, we find 48/543 (8.8%) traits with evidence of colocalization (CLPP>=0.01), comprising 58 unique outlier-associated variants.

## Discussion

Molecular QTL in pluripotent cells provide insights into the spectrum of genetic effects with trait-relevance that can manifest during development and across cell differentiation. To maximize the power to identify such QTL, we harmonized existing population-scale iPSC genetic and transcriptome datasets across five studies, and generated additional data. The scale of our resource, spanning iPSCs from close to one thousand individuals, has enabled the generation of a comprehensive map of *cis*-QTL across a broad range of expression phenotypes, but also *trans*-eQTL and rare variant effects that are typically power-limited in smaller studies.

We identified *cis*-QTL for each of the five gene phenotypes we considered in our genetic analysis, yielding *cis*-QTL for a total of 58% of all quantified genes (N=18,556). This includes 994 *cis*-eGenes that were not previously detected in other tissues and 3,118 *cis*-QTL genes with QTL that are specific to ratio- or exon-level QTL effects. Next to the *cis*-QTL, we identified 193 *trans*-eQTL; 24% of the *trans*-eQTL were driven by ratio- or exon-level *cis*-eVariants, variants that are commonly omitted in *trans*-eQTL analyses. Even when a gene-level effect was observed for a *trans*-eVariant, we observe that 91 (out of 139) of these *trans*-eQTL are also linked to ratio *cis*-eQTL. This demonstrates the existence of *trans* effects downstream of *cis*-QTL that do not affect gene expression level but instead act on other gene expression phenotypes.

To assess the impact of rare variants, we analyzed their role in gene expression outliers. Similar to observations in previous studies in somatic tissues^39^, rare SNP, indel and SV variants were enriched in outliers compared to non-outliers. By comparing iPSC against 35 GTEx tissues, we observed that iPSC provides the greatest power for detecting aberrant gene expression associated with rare, highly-conserved SNPs. Furthermore, when looking at genes where outlier effects were detected in iPSC but not in any GTEx tissue, we find a unique subset of genes important to developmental processes and linked in OMIM to rare developmental disorders, such as the gene *PRSS12* (mental retardation)^40^, *RASGRP2* (bleeding disorder)^41^ and *DUOX2* (thyroid dyshormonogenesis)^42^.

The large-scale QTL maps on the diverse set of gene expression phenotypes made it possible to generate an extensive colocalization map between the QTL effects and GWAS loci. We annotated over 4,400 GWAS implicated loci (out of the 29,666 assessed loci), originating from over 600 traits, to expression changes in iPSCs. We observe unique colocalized loci to each of the *cis*-QTL types and a wide range of traits, from physical traits to diseases and lab-measurements. Over 40% of the GWAS colocalized QTL loci are found not to be linked to gene-level eQTLs, highlighting the importance of effects beyond gene expression level. Interestingly, we also observed colocalizations for two of the *trans*-eQTL hotspots; namely, the hotspots around *ELF2* and *CREB3L2*, which both colocalized with height GWAS signals. Furthermore, we found colocalization for developmental traits, such as congenital craniofacial abnormalities, and cancers. Lastly, by integrating our colocalization results and rare variants linked to expression outliers, we demonstrated prioritization of variants with large impacts on phenotype in the UK Biobank, thereby providing a means for understanding the spectrum of large-effect rare variants associated with a wide range of complex traits and diseases.

Overall, we present the most comprehensive map of genetic regulation of gene-expression in iPSC to date, and demonstrate the use of this map by generating large-scale colocalization between GWAS and iPSC QTL. Here, we were able to link both the QTL map as well as the outlier linked rare variants to trait-associated genetic variants, and observe relevance for developmental traits, physical traits and cancer, among multiple others. Large-scale characterization of iPSC presented herein, combined with population-scale cohorts on differentiated tissues previously published^2,4,5^, allows for detailed comparisons of the molecular functions of genetic variants throughout different stages, from a pluripotent to terminally-differentiated tissues. These complementary datasets, will be invaluable in ongoing research efforts such as gene therapies, by showing a more complete context in which genetic variants and genes function.

The genetic maps and colocalization catalogs generated by the i2QTL consortium form a unique reference catalog, further aiding interpretation of risk variants in a unique cell type relevant for both development, cellular differentiation and cancer research. We expect that the genetic maps presented here, in ongoing combination with GWAS results will reveal the molecular underpinnings of genetic disease and traits manifesting during early-development.

## Online Methods

### Dataset information

Within the “Integrated iPSC QTL” (i2QTL) consortium we reanalyzed iPSC data originating from five different studies. Below a short description is given on each of the analyzed studies, in **table S1** the references to the data sources are given.

#### HipSci

The Human Induced Pluripotent Stem Cells Initiative (HipSci)^9,43^ was set up to generate a large, high-quality reference panel of human iPSC lines for the research community. Within the initiative a large population based iPSC panel was formed, next the population based inclusion there was a special inclusion for several diseases: Monogenic diabetes, Bardet-Biedl syndrome, hereditary cerebellar ataxia, hereditary spastic paraplegia, kabyki syndrome, usher syndrome and congenital eye defects, congenital hyperinsulinia, alport syndrome, hypertrophic cardiomyopathy, primary immune deficiency, bleeding and platelet disorders, macular dystrophy, retinitis pigmentosa, batten disease and childhood nerolog diseases. iPSC lines were generated through a non-integrative methodology (Sendai virus) using either fibroblasts or blood as a starting population. All of the lines were genotyped using the Illumina beadchip HumanCoreExome-12 genotyping chip. RNA-sequencing was performed, using either a pair-end stranded protocol, or a single end protocol, followed by Illumina sequencing. For a subset of the samples whole genome sequencing data was generated. Next to transcriptomic and expression data DNA-methylation information was generated using the Illumina 450K or Illumina EPIC array. For QC information on embryonic stem cells and the donor material (i.e. fibroblast or blood), was generated on a subset of the lines. In this study, we considered data from N=543 donors. A subset of (N=166) has previously been described in Kilpinen et al. More information about data generation, which is the same for the entire consortium, can be found in the original publications Kilpinen et al and Streeter et al.

#### iPSCORE

The iPSCORE project^6,44,45^ was completed with the goal of assembling high quality iPSC lines derived from hundreds of individuals, and profiling them with an array of genomics assays, to provide a resource for studying functional genetics in the context of derived human cell lines. For 273 iPSCORE individuals, deep whole genome sequencing (median 48x) was performed and additionally 215 of these individuals had iPSCs created by reprogramming fibroblasts, on which RNA-sequencing and chip genotyping using the Illumina MEGA array was also performed. Notably, some iPSCORE individuals are related as part of families of 2-14 subjects, while 167 are unrelated. Further information about this study, its constituents, and the generation of the sequencing data can be found in several previous publications (Panopoulos et al., DeBoever et al., D’Antonio et al,).

#### GENESiPS

The original GENESiPS study included 201 subjects with insulin sensitivity measurement performed by a modified insulin suppression test in accordance with Knowles et al^46^. The aim of the study was to generate an iPSC library reflecting the broad spectrum of insulin sensitivity in human populations. iPSC lines were generated through a non-integrative methodology (Sendai virus) using erythroblasts as starting population. iPSC grown under feeder-free conditions from passage 8-11 were used for RNA-sequencing on the Illumina HiSeq 2500 system with 100 nucleotide single-end reads. Further information can be found at Carcamo-Orive et al^10^.

#### PhLiPS

The PhLiPS Study (Phenotyping Lipid traits in iPS-derived hepatocytes Study) aimed to create a library of iPSC lines and iPSC-derived hepatocytes of diverse genotypes for metabolic profiling and lipid trait genetic screening. Detailed methods and data descriptions can be found elsewhere Pashos et al^7^. As a part of the Next Generation Genetic Association Studies (Next Gen) program, PhLiPS ascertained 91 subjects who were free of cardiovascular disease and in generally good health. Peripheral blood samples obtained from the subjects were used for genome-wide genotyping, blood lipid measurements and for generating iPSC lines. Infinium Human CoreExome-24 BeadChip (Illumina) was used for all sample genotyping. Extracted RNA with a minimum RNA integrity number (RIN) 7.5 were sequenced on HiSeq 2000/2500 systems (Illumina) with paired-end, 100-bp/125-bp read lengths with a target read-depth of 50 million reads per sample.

#### Banovich

The Banovich et al study^8^ investigated the use of iPSCs to study the impact of genetic variation on gene regulation. A panel of iPSC lines were derived from 58 Yoruba (YR) lymphoblastoid cell lines (LCLs), originally generated from the YRI samples within the 1000 Genomes (1000G) projects. LCLs were reprogrammed using an episomal approach described in Okia et al^47^. RNA-sequencing was performed using fifty-base pair single-end libraries using the Illumina TruSeq kit, sequenced on an Illumina HiSeq 2500. More information on the iPSC generation and RNA-sequencing can be found in Banovich et al. Genotypes were taken from the 1000G samples^48^ where the LCLs are originally derived from, for this we focused on the array genotypes to match the other datasets.

### Genotyping information

#### Array based genotyping

Genotyping information for the individual studies was generated as described in the original publications referenced above. For the QTL analyses the genotyping information as generated by genotyping arrays are used and human genome assembly GRCh37 was used. The genotype information was imputed and phased per sample using IMPUTE2 v2.3.1^49^ for imputation and SHAPEIT2 v2.r790^50^ for phasing. The imputation and phasing was performed using a combined genotyping reference, encompassing the haplotypes from the UK10K cohorts and (1000G) Phase 1 data^50,51^. The imputation was run in chunks of average 5 Mb and 300 kb buffer regions on each side, and used the following MCMC options (-Ne 20000 -k 80) for autosomes. SHAPEIT2 was run without MCMC iteration (-no-mcmc) so that each sample is phased independently. Single-sample VCFs were merged and subsequent QC was performed using Genotype Harmonizer^52^ and BCFtools^53^. After imputation each cohort was combined and cohort-, and therefore chip-, wise SNP-QC was performed using Genotype Harmonizer. SNPs were excluded if the call rate was below 90% and imputation quality (Mach R2) was below 0.4. After dataset wide QC we merged the datasets and performed a combined SNP QC filtering for a call rate of 1.0 and an imputation score of 0.4. Finally we were left with 2,533 samples and 7,188,631 variants for which array genotypes were available.

#### Super population assignment

All imputed samples were assigned to 1000G super-populations by projecting the samples with the genetic PCA based on the full 1000G reference (phase 3) dataset^48^. Next we selected the variants that had a minor allele frequency (MAF) over 5% within 1000G, were available within the imputed i2QTL genotype dataset and were retained after pruning of the remaining variants. We performed a principal component analysis (PCA) on the 1000G samples and projected the i2QTL samples on to the calculated PCs. Next we calculated the distance for each of the i2QTL samples to the centroid value for each super-population as calculated using the 1000G reference, based on the first 40 principal components (**Figure S8**). We chose to use 40 components to predict the super-population, as this was the optimal number based on re-assigning the 1000G samples back to their original super-populations.

#### Whole genome sequencing

For a subset of the samples, originating from HipSci and iPSCORE, whole genome sequencing data was available. Within the HipSci consortium whole genome sequencing data was available from fibroblasts obtained from 204 donors, many of which also had whole genomes available from derived iPSCs. From the iPSCORE consortium, 273 individuals had whole genome sequencing data available which was generated from blood (254 donors) or fibroblasts (19 donors). Genotype calling on the i2QTL WGS samples is described in detail in Jakubosky et al^54^. In short, we called both SNVs and structural variants within the joint dataset. To do so we: (1) called SNPs and short indels using GATK’s best practices for genotype calling; (2) called duplications, deletions, inversions, reference mobile element insertions (rMEI) and other novel adjacencies referred to as “breakends” using SpeedSeq (LUMPY/CNVnator); (3) called duplications, deletions and multiallelic copy number variants (mCNVs) using Genome STRiP; and (4) called mobile element insertions using MELT. After calling the structural variant classes, subsequent quality control and a stitching and merging procedure was used to derive a non-redundant high quality set of variants. In the outlier analyses a total of 34,804,470 SNPs and 4,721,392 indels were called in 928 high-quality samples. 528 of these samples had iPSC RNA-seq.

### RNA-sequencing

#### RNA-sequencing feature quantification

Raw RNA-seq data for the samples of each of the projects, including Choi et al^13^ as reference, was collected from EGA, dbGaP and SRA (**Table S1**). Depending on the availability of data from each source, we started the reprocessing with CRAM, BAM or FASTQ files. To ensure uniform processing of all samples, the CRAM and BAM files were first transformed to FASTQ files. The reads in the FASTQ files were trimmed to remove adapters and low quality bases (using Trim Galore!^55–57^). Next, trimmed reads were alignment using STAR (version: 020201)^58^, using the two-pass alignment mode and the default parameters as proposed by ENCODE (c.f. STAR manual). All alignments were relative to the GRCh37 reference genome and Ensembl 75 genome annotations^59^. Based on the quality controlled aligned reads we quantified gene-level information on all samples. For the paired-end stranded data we also quantified transcript-ratios, exon-level, alternative polyadenylation ratios and splicing-ratios.

Gene-level RNA expression was quantified from the STAR alignments using featureCounts (v1.6.0)^60^, which was applied to the primary alignments using the “-B” and “-C” options in stranded mode if applicable, using the Ensembl 75 GTF file. In case multiple RNA-seq runs per iPSC line were generated these were summed to one set of gene-counts per iPSC line. Read counts per iPSC line were normalized for gene length and the edgeR^61^ normalized total expression count per sample, yielding corrected transcript per million counts (TPM). Using the same setup exon expression level were quantified. We chose to first do a strand-specific merge of overlapping exons and quantified these meta-exons as one entity. Subsequent normalization is the same as for the feature-count derived gene information.

To facilitate a comparison to GTEx v7^5^ a second gene-level quantification was performed. The STAR aligned BAMs were used to quantify gene-level information using RNA-SeQC^62^ (v.1.1.8) with the gencode v19 annotation, matched to the GTEx annotation. This limited the quantification differences to only read-QC and alignment. This RNA-SeQC quantification was used only when comparing the i2QTL data to GTEx tissues.

Transcript ratios were quantified by determining transcript isoform levels using Salmon^63^ (version: 0.8.2). The Salmon transcript database was built based on transcript information from Ensembl 75. Salmon directly uses the QCed FASTQ data; in the transcript quantification “--seqBias”, “--gcBias” and “VBOpt” options were used. The transcript count quantification as returned by Salmon were normalized as described for the gene-level quantification using feature count information. Subsequently, we transformed the transcript levels to ratios per gene, by dividing the count of an individual transcript by the transcript sum per gene.

Alternative polyadenylation (APA) quantification was performed following a similar approach to that used in Zhernakova et al^4^. In brief: first, overlapping 3’UTRs for transcript isoforms from the same gene were merged by taking the union of overlapping regions, such that the set of 3’UTRs for each gene was a set of disjoint genomic regions. Next, these regions were extended by considering the most distal available polyadenylation site for each 3’UTR region, as present in the APADB^64^. For each 3’UTR, the set of alternative polyadenylation sites within that region were identified by taking the union of all annotated APA sites from the APADB, in addition to the annotated end of the 3’UTR given by the Ensembl annotation. These APA sites then subdivide each 3’UTR into a set of windows. The depth of RNA-seq in each window was then computed by using the samtools bedcov function. Finally, the read depth in each window was expressed as a fraction of read depth in the window immediately upstream.

Lastly, we quantified splicing levels using leafcutter^65^ (v0.2.8). Quantifications were of splice-events are calculated as described in the leafcutter manual. First the junction reads are extracted from the BAM files, after which leafcutter clusters junction reads into introns. Subsequently the intron counts are transformed into ratios per cluster and we linked the intron locations to genes.

#### RNA quality control

After feature quantification low quality RNA-seq samples were filtered out by applying filters on both Picard^66^ and VerifyBamID^67^ quality measures as well as gene expression levels. We defined high quality samples as those with > 15 million reads, > 30% coding bases, > 65% coding mRNA bases, a duplication rate lower than 75%, Median 5‘ bias below 0.4, a 3’ bias below 4, a 5’ to 3’ bias between 0.2 and 2, a median coefficient of variation of coverage of the 1000 most expressed genes below 0.8, a free-mix value below 0.05. Lastly, we removed samples that had low expression correlations (<0.6) to the average iPSC expression values across our study, as measured per chromosome. This resulted in 1,367 iPSC lines derived from 948 donors for analysis, all of which also have genetic information available. Next to this we had 98 samples from Choi et al^13^ and HipSci^9,43^ which are included as reference (table S2).

After RNA QC the sample linking was validated using VerifyBamID, for each of the RNA-samples the best matching genotype were computed and compared to the expected match. This resulted in the identification of 36 sample mixups (between 0-26 mismatches per study), and identified in total 33 unmatched RNA-seq samples. Some of the mix-ups were corrected (N=34) and others were removed (N=2).

#### PEER correction and optimization

We correct for transcriptome-wide confounders arising from the merging of multiple datasets, to do so we chose to estimate PEER^21^ factors to capture these confounders. We chose not to include any known variation when estimating PEER factors, as meta-data was sparse and not standardized between studies. We ran PEER (v1.3) on the log-transformed gene-level quantifications, on genes that had a TPM > 2. We assessed the impact of the number of PEER factors on eQTL mapping in our data by calculating the number of eGenes (at least one eQTL at FDR < 5%) with incrementally increasing number of PEER factors (**Figure S9**). We chose to use 50 PEER factors to correct for confounders in out data. Assuming *trans*-eQTL impacting many transcripts may be included as latent factors in PEER analyses, we then tested eQTL effects against each of the 50 PEER factors. We did not find any likely *trans*-eQTL-associated PEER factors.

### Quantitative trait loci mapping

For the quantitative trait loci mapping (QTL) mapping, both the *cis* and *trans*, we used a linear mixed model implemented in LIMIX^68^. This model allowed to control for both population structure and repeat lines from the same donor using kinship as a random effect component. The kinship matrix was estimated using the identity-by-descent function in PLINK (1.07)^69^, on independent variants with a MAF >5%. Fifty PEER factors, derived on the gene-level data were included as fixed effect covariates in all analyses, optimization is described above. The QTL mappings are performed on the log transformed standardized expression levels in case of the gene-level and exon-level data; for the other expression phenotypes, the ratio based phenotypes, we used an arcsin-transformation.

To control for multiple testing we used an approximate permutation scheme as presented in Fast-QTL^70^, based on a parametric fit to the null distribution, to adjust for multiple testing across *cis* variants for each gene. Briefly, for each gene, we obtained p-values from 1,000 permutations on the genotypes while keeping covariates, kinship, and expression values fixed. Subsequently, we estimated an empirical null distribution by fitting a parametric Beta distribution to top P-values per gene per permutation round. Using this null model, we estimated *cis* region adjusted p-values for QTL lead variants. To control for the multiple genes-level tests we do per QTL run, we performed the Storey’s Q-value procedure^71^. In cases where multiple features per gene where tested, i.e. for transcript-ratio, exon-level, splicing-ratio and APA-ratio *cis*-QTL (herein features), we chose to control the FDR at gene-level, to explicitly account for the number of features per gene. To do so, we applied a Bonferroni correction, for the number of features per gene, on the permutation-based *cis*-SNP corrected eQTL p-values. We then take the lead QTL effect per gene, i.e. the most significant associated variant-feature pair (*cis* region and across features adjusted), and apply the Storey procedure on these selected p-values. After the Storey procedure we find the Q-value (FDR) threshold and report all QTL at the selected level, i.e. potentially more per gene, that pass this threshold.

#### *cis*-eQTL Mapping

During the mapping of *cis*-quantitative trait loci (*cis*-QTL), we considered common variants (MAF>1%) in gene-proximal regions of 250k upstream and downstream of gene transcription start and end sites (GRCh37). We limited the analysis to include only paired-end stranded European samples, as determined from the super population assignment, (n=716 donors, n=932 lines) to limit variation between trait discovery and to have the same discovery sample size in all *cis*-QTL maps. We controlled for multiple testing at a gene-level FDR of < 5% for the *cis*-QTL maps.

For the different expression phenotypes, we tried to use matched inclusion criteria. We limited the analysis to features that were expressed, i.e. non-zero expression, in at least 25% of the samples. For the splicing and APA-QTL we required at least 50% of the samples to have a non-zero and non-NA ratio, the assessed genes per QTL type is summarized in **Table S15**.

After identification of the lead *cis*-eQTL for the gene-level effects, we searched for independent eQTL (secondary eQTL and higher order effects). For each of the significant eQTL gene the significant eQTL variant(s) from lower levels were added as a covariate to the model and a new QTL mapping round was run to identify independent *cis*-QTL. This search for independent effects was continued for each gene until no more independent *cis*-eQTL were identified.

#### Trans-eQTL mapping

During the mapping of *trans*-quantitative trait loci (*trans-e*QTL), we considered common variants (MAF>1%) in gene-distal regions defined as at least 2.5M upstream and downstream of the gene transcription start and end sites (GRCh37). Given the large number of tests in a full *trans*-map, we chose to limit the *trans*-eQTL map to both *cis*-QTL variants and GWAS implicated variants. This limited the number of variants that had to be tested to 115,709. In addition, we only considered protein coding genes with a TPM ≥ 1 (n=17,039) and expressed in at least 25% of the samples. Other settings were matched to those described above. However, to limit the chance of spurious associations we included an extra expression normalization step - we quantile normalized the normalized gene counts to match a normal distribution and we regressed out the *cis*-eQTL variants identified before. During the *trans*-QTL phase we switched to using all samples with European ancestry (n=743, lines=1,120) as we only looked at gene-level effects.

To further limit the chance of spurious associations, specifically those linked to cross-mapping of RNA-reads^72^, we set out to black-list combinations of genes a-priori. We created a black-list by using the information contained in the mapping of the RNA reads to the genome, i.e. using primary and secondary genome matches. The gene-level quantifications are based only on the primary mapping of a read but the secondary mappings that do pass the mapping thresholds are informative to identify genes with high sequence similarity. A benefit of this approach is that it is directly based on the reads that we also use for gene quantification and is influenced by the same biases. We built this cross-mapping map using all the paired end stranded data. When a gene combination would be in the black list we also removed the *trans*-window around the secondary gene from the test region. We also excluded the HLA region, due to its complex LD structure.

### *cis*-eQTL annotation

The i2QTL genetic maps where annotated by overlapping the QTL signals with information taken from published QTL maps, gathered from the original iPSC studies^7–10,45^, GTEx^5^ and BIOS^4^.

To assess the replication of the eQTL effects published in the original iPSC studies^7–10,45^, we tested for replication of the published eQTL effects at p-value<0.05 and assessed effect direction **(Fig. S1-4)**. The original QTL maps were taken as provided, i.e. replication was assessed based on the published significance levels. When assessing the replication within the iPSCORE study we focused only on the effects attributed to SNPs, and excluded the effects of structural variants. Between 75% and 82% of the effects were assessed in both the original study and the i2QTL study, replication rates of the effects that are between 62% and 86% **(Fig. S1-4)**.

Using the GTEx^5^ and BIOS^4^ QTL maps we assessed the overlap of eGenes to identify effects specific to iPSC. Effects from the original studies where taken as provided, i.e. FDR 5% as calculated in the studies. Next we assessed replication in terms of eGenes within the individual tissues (**Fig 1B)** as well as looking at eGenes in any of the GTEx tissues or BIOS (**Fig 1B-1C**).

To test for similarity between genetic effects observed at the different gene expression phenotypes, we assessed replication between the signals at the different levels. The overlap was assessed for individual eQTL, i.e. SNP-gene combinations, by mapping the different QTL types to the respective gene of the effect and taking the most significant association (at gene-level FDR) if multiple were tested per gene. If the effect present at the first QTL type was replicated at FDR 10% in a second QTL type this was counted as a replicating effect (**Fig 1E**).

### Replication of *trans*-eQTL

Using the expression samples from other inferred super populations (n=186 donors, n=253 lines) **Table S2**, we could assess replication of the *trans*-eQTL. Next to this we used DNA-methylation data available in a subset (n=572 donors, n=841 lines) to do cross-omic replication of the identified *trans*-eQTL.

The replication of the *trans*-eQTL in the left out expression data was performed using the same setup as for the main *trans*-eQTL analysis, with the exception of the PEER factor correction. To not induce effects based on PEER factors, we chose to only correct for the limited known factors across our datasets: inferred ancestry, sequencing type (paired-end (yes/no), stranded (yes/no)) and hot encoded vectors describing the dataset of origin. We deemed *trans*-eQTL as replicated in the left-out samples when the raw p-value of the association was below 0.05 & the effect direction was matching to the effect observed in the *trans*-eQTL map.

The replication within the DNA-methylation data was similarly matched as closely as possible. We started with a joint normalization of the Banovich et al^8^ and HipSci^9,43^ DNA-methylation arrays. As the data was generated on two different Illumina methylation arrays, the Illumina 450K and Illumina EPIC array, we started with sub-selecting the CpG probes that are present on both. After this we normalized the DNA-methylation profiles, as described in detail Bonder et al^3^, based on the DASEN^73^ normalization. After joint normalization, we correct the data for the first 20 PCs to account for batch effects in the DNA-methylation data. In order to replicate the *trans*-eQTL in DNA-methylation data, we first had to link CpG probes to genes. We chose to do so by linking a gene to all DNA-methylation probes inside the gene and CpG probes that are within within 250Kb of the gene TSS and TES. Given that we test a multitude of CpG’s per *trans*-eQTL gene we determine the number of independent DNA-methylation probes per gene (R2<0.2) and used Bonferroni procedure to correct for the number of independent probes that were tested per gene. We deemed *trans*-eQTL as replicated in DNA-methylation when the CpG corrected replication p-value was below 0.05.

To check if we replicate more effects than expected by chance, we repeated the *trans*-eQTL and *trans*-meQTL replication on random *trans*-eQTL pairs. To try and match the trans-eQTL characteristics, we chose to link the *trans*-eQTL variant to a gene with matched expression characteristics, by selecting the ten genes per *trans*-gene with the closest average and variance levels. The same random pairs were used for the *trans*-meQTL replication.

### Outlier analysis

In addition to variation linked to common variants (MAF >1%), rare variants were linked to transcriptomic outliers. Using the featureCount gene quantifications (log TPM), genes were subset to only include autosomal protein-coding and long non-coding RNA genes. Cell lines from donors with predicted ancestry other than European super-population were removed, and only paired-end stranded cell lines were selected. Genes were then filtered for minimal expression, defined as gene expression TPM>0 in 50% or more within each study.

To correct for technical effects, PEER^21^ factor correction was run on the filtered data as described above. Following the correction of 50 PEER factors, as chosen based on the *cis*-eQTL PEER optimization, the resulting residuals were scaled and centered (Z-score). Samples were then checked for large gene outlier counts (i.e. cell line is found to be the most under- or over-expressed cell line across hundreds of genes), which suggests possibly incomplete correction for technical effects. Cell lines with expression abs(Z-score)>2 in more than 100 genes were removed from subsequent analysis (N=21 lines). Finally, cell lines were retained only if WGS was available in addition to RNA-seq, leaving data from the HipSci and iPSCORE projects only. After applying these steps, 17,514 genes and 425 cell lines remained in the dataset for further analysis.

To prepare the WGS genotype data SNP and indel variants for the analysis, variants were first filtered based on QVSR tranche 99%. The software vcfanno^74^ was used to annotate the WGS VCF with minor allele frequency from gnomAD^75^ (version r2.0.2), and CADD score from CADD^76^ (version 1.3). Variants were filtered per sample to retain variants with at least one alternate allele only. Variants were then linked to genes using the bcftools^53^ window command, with a maximum distance of 10 Kb window size and using the Ensembl 75 GTF as reference. A separate file was produced for each cell line comprised of the following columns: cell line ID; gene ID; chromosome; position; gnomAD MAF; CADD (phred); CADD (raw).

We performed a comparative analysis using GTEx v7 tissues, reprocessing i2QTL data to match the GTEx v7 processing pipeline to limit technical variation. For this specific analysis, RNA-SeQC expression quantification was used, and a separate PEER analysis was run to correct for technical variation, without known factors. As before, the top 50 PEER factors were chosen for i2QTL data. For GTEx v7 tissues with <= 150 samples, 15 PEER factors were used; for tissues with <= 250 samples, 30 PEER factors; for tissues with >250 samples, 35 PEER factors. GTEx v7 WGS variants were annotated with MAF from gnomAD (version r2.0.2) and CADD scores from CADD (version 1.3) using vcfanno. GTEx v7 tissue samples with expression abs(Z)>2 in more than 100 genes were removed. Tissues were retained only if there was >=50 samples available (N tissues = 35). Residuals resulting from PEER factor correction were centered and scaled to generate expression Z-scores.

#### Outlier enrichment

An enrichment score was calculated to quantify the difference in proportions of outlier lines with variants across several MAF/CADD bins compared with non-outlier lines. Specifically, enrichment here refers to the relative risk:

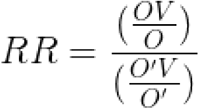
where *OV*is the number of outlier lines with >=1 variant in or near (+/− 10 Kb upstream or downstream of gene body) a gene passing given MAF and CADD thresholds, *O* is the total number of outlier lines, *O’V* is the number of non-outlier lines with >=1 variant in/near gene passing given MAF and CADD thresholds, and *O*’ is the number of non-outlier lines. The relative risk is reported with 95% Wald confidence intervals derived from the asymptotic distribution of the log relative risk:

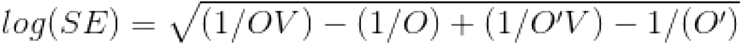

The bounds on the confidence interval are defined as follows:

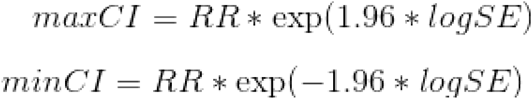

The analysis was performed separately for SNP, indel, SV variants, and across different MAF bins (from common to rare) and CADD bins (progressively more deleterious variants). Expression outlier direction (i.e. under-expression, over-expression) was tested separately. For example, for under-expression outliers, an outlier line was defined as the least-expressed line in a given gene that also has a Z-score < −2. In this way, a gene can have a maximum of one outlier line per outlier direction. Non-outliers were defined as lines with a Z-score between −1 and 1 for a given gene. Genes were discarded if there was not at least one outlier and one non-outlier line matching MAF and CADD thresholds. The outlier analysis was performed for both i2QTL and GTEx datasets.

### Colocalization of GWAS loci with iPS QTL

To choose the combinations of loci, features, and traits at which to test for colocalization, we started with two sets of curated GWAS summary stats: one including UK Biobank (UKBB) rapid GWAS results (for 1,740 traits) (cite Neale et al.), and the other including publicly available GWAS results gathered from the NHGRI-EBI GWAS catalog^22^ and PHENOMESCANNER V2^34^ for 303 studies, obtained through a wide variety of studies and consortia^35^. For every trait in this set, we selected loci having a GWAS association a p-value of < 5×10^−8 and located at least 1Mb away from all previously selected (more significant) loci for the same GWAS trait. We further narrowed this set of GWAS-relevant loci to with a significant *cis*-QTL, for any of the quantified expression phenotypes, within 10kb of the lead GWAS hit at a gene-level corrected FDR of less than 5%. Due to the vast number of SNPs and traits in the UKBB, we only tested UKBB GWAS hits for colocalization if the lead GWAS hit overlapped exactly with a known eQTL, for computational feasibility. The resulting loci comprised our colocalization test set, to be tested for colocalization with every eQTL phenotype having a significant association nearby. Given the presence of several different types of eQTL and abundant measured features for each of these eQTL types, it was possible for a single GWAS locus to be tested for colocalization with a number of QTL traits originating from single or multiple genes.

We then tested every resulting pair of GWAS locus and eQTL feature in our set. For each of these pairs, we narrowed our summary statistics to the set of SNPs tested for association with both the given GWAS trait and the given eQTL trait, and removed all sites containing less than 5 SNPs after this filter. Using the full 1000 Genomes dataset from phase 3 (2,504 individuals) as a reference population^48^, we estimated LD between all of these SNP pairs. We then ran FINEMAP^32^ independently on the GWAS and eQTL summary stats to obtain posterior probabilities of causality for each of the remaining SNPs, and we combined these probabilities to compute a colocalization posterior probability (CLPP) using the formula described in the eCAVIAR method^32,33,35^.

Given the imperfect overlap between the variants in the GWAS summary statistics and the QTL data it can happen that the minimal corrected eQTL p-value or the GWAS p-values in a locus, after subsetting, were higher than the initial inclusion criteria of a locus. To exclude colocalizations in loci where only a very weak association, at either or both statistics, is observed, we excluded tests where the minimal GWAS p-value in the overlapping locus was higher than 5e-6 or the gene-level corrected eQTL p-value was greater than 0.05. We considered a locus to be colocalized if it passed one of three CLPP thresholds; 1) minimal 5 variants with a CLPP of 0.5; 2) minimal 10 variants and a CLPP of 0.1; or 3) minimal 25 variants with a CLPP of 0.01.

Last, to also link *trans*-eQTL to GWAS loci we performed an extended *trans*-eQTL mapping linking all variants in 250Kb around the identified *trans*-eQTL variants to the identified downstream genes. This trans-eQTL information was subsequently used to perform a colocalization analysis as detailed above for *cis*-QTL.

### Annotating i2QTL rare variants using UKBB GWAS summary statistics

To test for differences in outlier and non-outlier associated rare variants and risk for disease and traits, we overlapped i2QTL WGS variants (as described in Outlier Analysis above) with those measured or imputed in UKBB GWAS^77^. Specifically, we looked at variants with gnomAD MAF<1% and CADD>0. Multi-allelic variants were discarded. Variants were retained if they were observed in only one i2QTL individual, with the hypothesis being that these variants are driving observed outlier effects. From this list of unique variants (i.e. observed in only one i2QTL sample), outlier-associated variants were found separately for under-expression and over-expression outlier samples (therefore, there could be a maximum of two outlier samples per gene).

For each gene with >= 1 outlier sample, non-outlier-associated variants were chosen for each non-outlier sample if a variant had a CADD score within a range +/− 5 of outlier variants. A non-outlier sample was defined as a sample with expression Z-score > −1 and <1 for a given gene. If a non-outlier sample had a greater number of variants than the outlier sample, variants were randomly downsampled to match the number of outlier variants. If a non-outlier sample had a less or equal number of variants than the outlier sample, all variants were chosen for that sample. This process was performed separately for each gene and each outlier direction (i.e. under-expression, over-expression). The final list of variants was linked to UKBB GWAS effect size and p-values for each trait in UKBB GWAS Phase 1.

### Data availability

All data used in the study is available via SRA, dbGaP or ENA; the full data availability is provided in **Table S1**. **Table S2** provides sample description on the samples used in the study. Full summary statistics on the significant cis-QTLs can be found on https://zenodo.org/record/3470764 (doi:10.5281/zenodo.3470764).

## Acknowledgements

The authors would like to thank the HipSci, iPSCORE, GENSIPS and PhLiPS cohorts for their input into the study and (early) access to the data. This work was supported by: the EMBL Interdisciplinary Postdoc (EI3POD) program under Marie Skłodowska-Curie Actions COFUND (grant number 664726) (to MJB & DS); National Institutes of Health (T32 LM012409 to CS; T15 LM01127 to DJ; U01 HL107388-01, P30DK116074, SPO 130829 to IC-O; HL107442, DK105541, DK112155 to KAF); Stanford Graduate Fellowship (to MJG); National Natural Science Foundation of China (31970554 to XL); EBI–Sanger Postdoctoral Fellowship (to NC); National Science Foundation Graduate Research Fellowship (to NMF); California Institute for Regenerative Medicine (GC1R-06673 to KAF). Research in the Stegle lab is supported by the BMBF, the Volkswagen Foundation and the EU (ERC project DECODE). SBM is supported by NIH grants U01HG009431, R01HL142015, R01HG008150 and U01HG009080. This work in-part used supercomputing resources provided by the Stanford Genetics Bioinformatics Service Center, supported by National Institutes of Health S10 Instrumentation Grant S10OD023452. The content is solely the responsibility of the authors and does not necessarily represent the official views of the National Institutes of Health.

## Author contributions

The main analyses and data preparations were performed by MJB and CS; DJ and MD performed structural variant calling and analysis; NMF performed GTEx v7 data processing; LF and XL performed rare variant annotation and interpretation; MJG performed the colocalization analysis; MJB, DS, BM and DH developed the QTL calling software; NC, IC-O, YP assisted with data processing and analysis; MJB, CS, MJG, SBM, OS wrote the manuscript; IC-O, NC, NMF, KAF, LF, MJG, DJ, BM assisted in editing the manuscript; MJB, CS, ES, KAF, OS, SBM conceived and oversaw the study.

