## Supplementary materials for "Systematic assessment of regulatory effects of human disease variants in pluripotent cells"

### Supplementary material

### Supplemental text 1: Trans GWAS findings

When assessing the effects of *trans*-eQTLs on GWAS we observed several interesting variants that overlap known biology. In the main text we highlighted several examples and additional notable *trans*-eQTL GWAS relations are described below. Furthermore, , tables S8, S9 and S11 contain all GWAS annotation of the *trans*-eQTLs. One of the four *trans*-eQTL hotspots was found to be directly linked to a GWAS variant for telomere length (rs412658), the hotspot on chromosome 19 around *ZNF257*. The GWAS variant is linked to eight genes, (*ZNF257*, *ZNF208*, *ZNF98*, *ZNF209P*, *RPL34P34*, *ZNF676*, *ZNF729* and *VN1R85P*) and in *cis* and seven *trans*-eGenes (*DNAH3*, *MATN4*, *RBPJL*, *GALNT13*, *BEST2*, *SLC30A8* and *S100A4*). *ZNF257* itself is significantly related to telomere length[1], however none of the downstream genes have previously been linked to telomere length. Another potential *trans*-eQTL of interest is the effect of a GWAS variant (rs2277339) for age of menopause to a *trans*-eQTL effect on *PRIM2*, the same eVariant is also linked to a *cis*-QTL on *PRIM1*. The effect of missense variants in *PRIM1* on age of menopause were already described in Stolk et al[2], however the effects on *PRIM2* are novel. The effects we identify are interesting as we identify a regulatory effect on *PRIM1* that is also directly linked to expression of *PRIM2*, *PRIM1* and *PRM2* function in a heterodimer at the protein level, but here we find a direct expression link between the two. Though interpretation of the *trans*- only eQTLs is more difficult,  we do find some interpetable signals, one of these effects being a breast cancer implicated variant (rs11199914); this variant is linked to the expression of *SGO2*. *SGO2* is part of the shugoshin gene family, which plays a role in chromosome stability in nuclear division[3]. *SGO1* part of the same family has already been shown to be involved in breast cancer[4].

Next to the GWAS variants being linked direct (or via LD), to *trans*-eQTLs, we applied colocalization around the *trans*-eQTL variants using the same approach as performed for *cis*-QTLs (see Results and Methods). The colocalization revealed several extra links between GWAS loci and *trans*-eQTLs results. We observed that two of the hotspots, the hotspots around *ELF2* and *CREB3L2*, are both linked to height[5] (**table S11**). Next to this we also find a hit for age-related macular degeneration[6] (rs10754220). A *cis*-QTL loci linked to a transcript ratio change of *TIRAP*, an adapter molecule associated with toll-like receptors, that has an effect in *trans* on the expression of *WNT6*. Both toll like receptors[7] and wnt signaling[8] has been implicated in age-related macular degeneration, however the direct connection of *TIRAP* and *WNT6* has not been described. Finally, a colocalization linked to the same genetic loci though a different lead SNP, we observe a *cis*-eQTL for *TIRAP* at gene level, that is linked to an expression change of *OSTN* in *trans* and is linked to LDL-levels by GWAS (rs11220462)[9]. *OSTN* (also known as *MUSCLIN*) was previously linked[10] to triglyceride levels such that this novel link to LDL-levels strengthens the metabolic role for *OSTN*.

### Supplementary figures

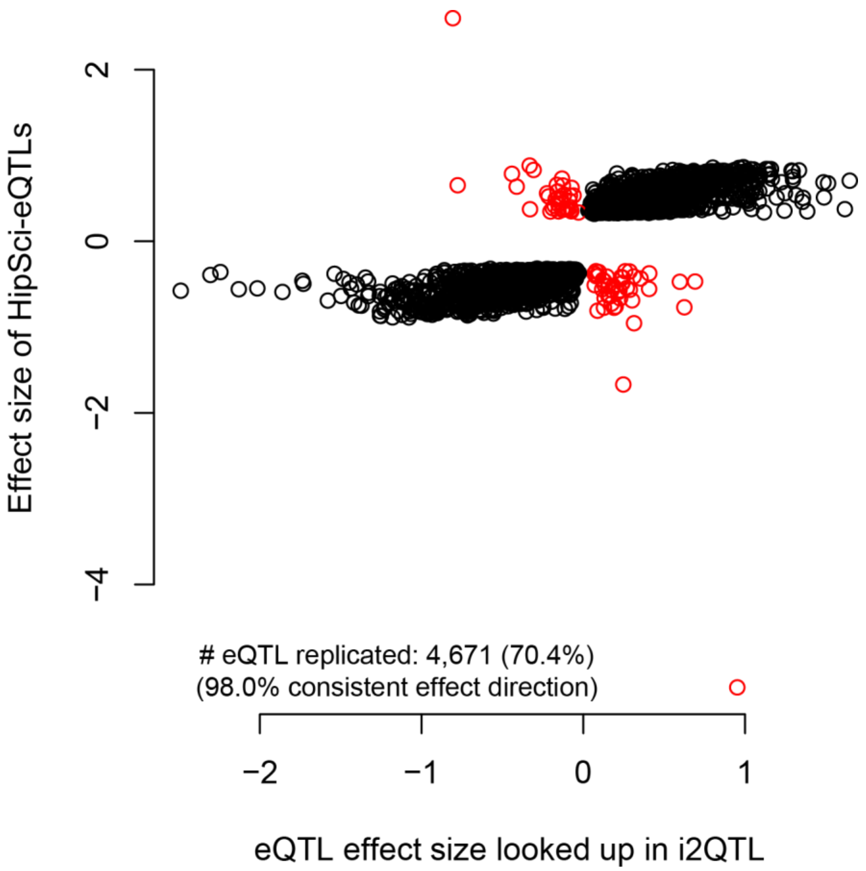


**Supplementary Figure 1.** Replication of the eQTLs effect sizes reported in HipSci Phase 1. On the Y-axis the original effect sizes as reported, on the X-axis within the i2QTL data. Over seventy percent of the effects are shared at a p-value cut-off at 0.05 within i2QTL and 98% of the effects are found to have the same effect direction.

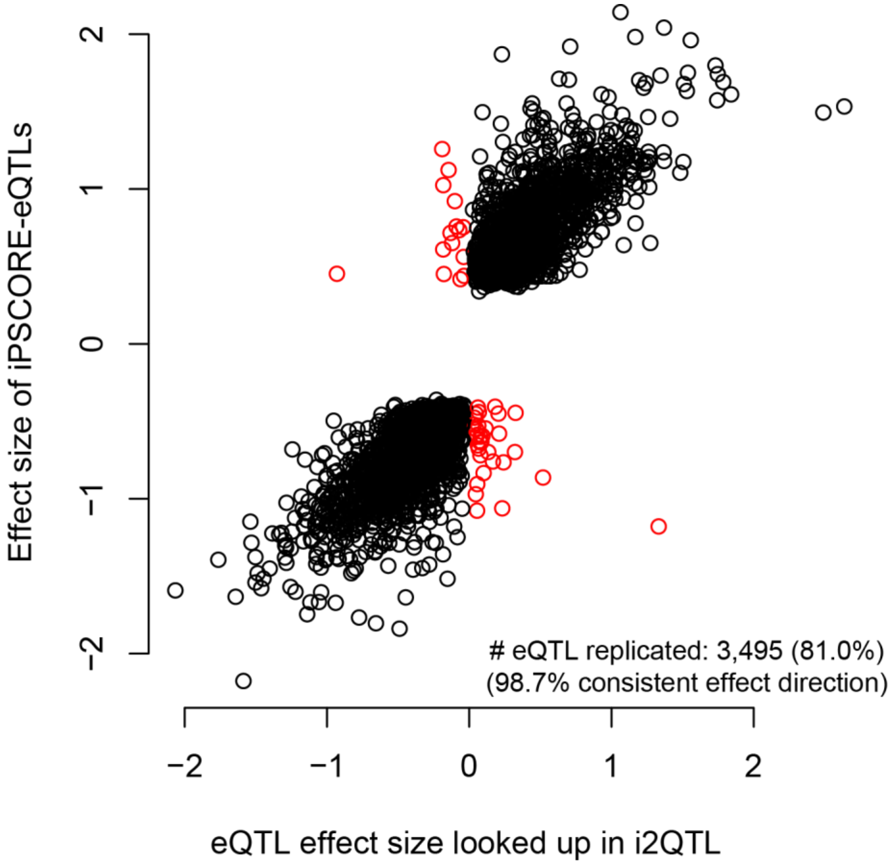


**Supplementary Figure 2.** Replication of the eQTLs effect sizes reported in the iPSCORE. On the Y-axis the original effect sizes as reported, on the X-axis within the i2QTL data. Over eighty percent of the effects are shared at a p-value cut-off at 0.05 within i2QTL and over 98% of the effects are found to have the same effect direction. Notable we only SNP effects, i.e. not the SV effects, are taken for replication.

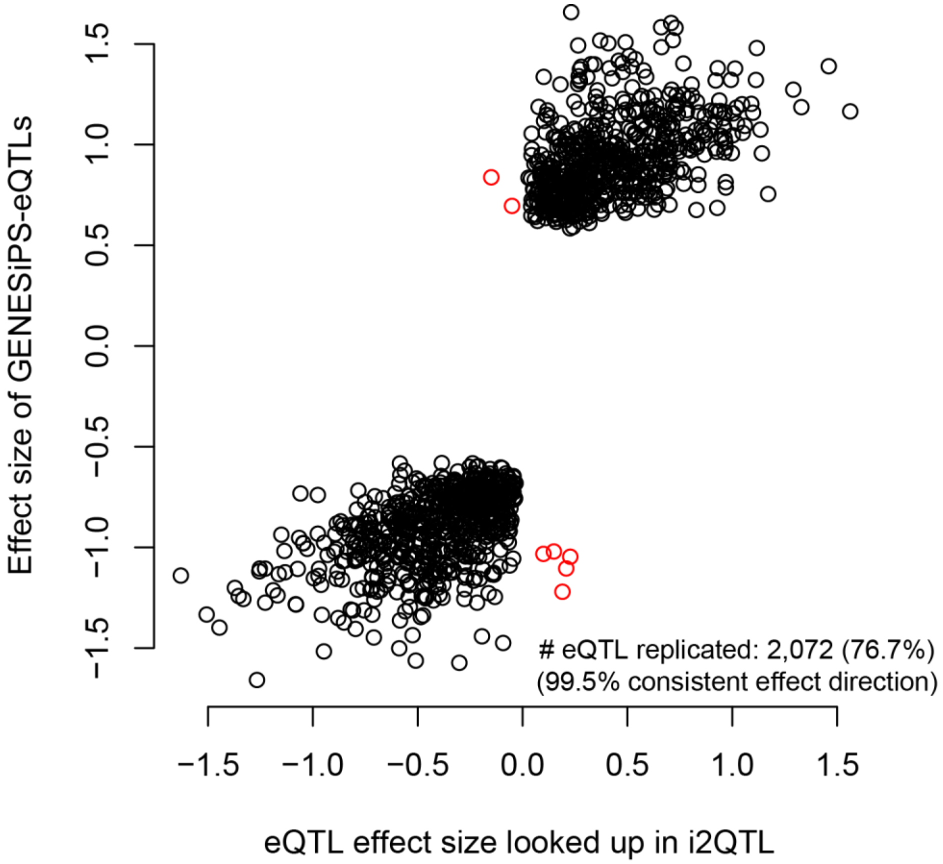


**Supplementary Figure 3.** Replication of the eQTLs effect sizes reported in the GENESiPS. On the Y-axis the original effect sizes as reported, on the X-axis within the i2QTL data. Over seventy percent of the effects are shared at a p-value cut-off at 0.05 within i2QTL and over 99% of the effects are found to have the same effect direction.

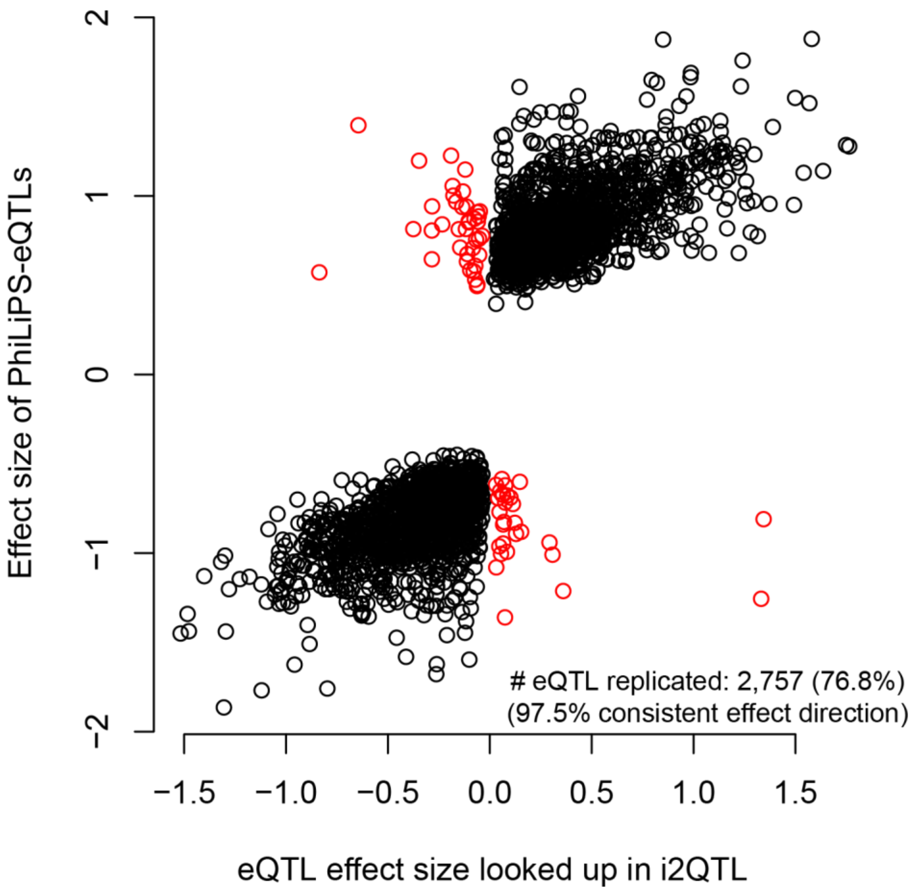


**Supplementary Figure 4.** Replication of the eQTLs effect sizes reported in the PhiLiPS. On the Y-axis the original effect sizes as reported, on the X-axis within the i2QTL data. Over seventy percent of the effects are shared at a p-value cut-off at 0.05 within i2QTL and over 97% of the effects are found to have the same effect direction.

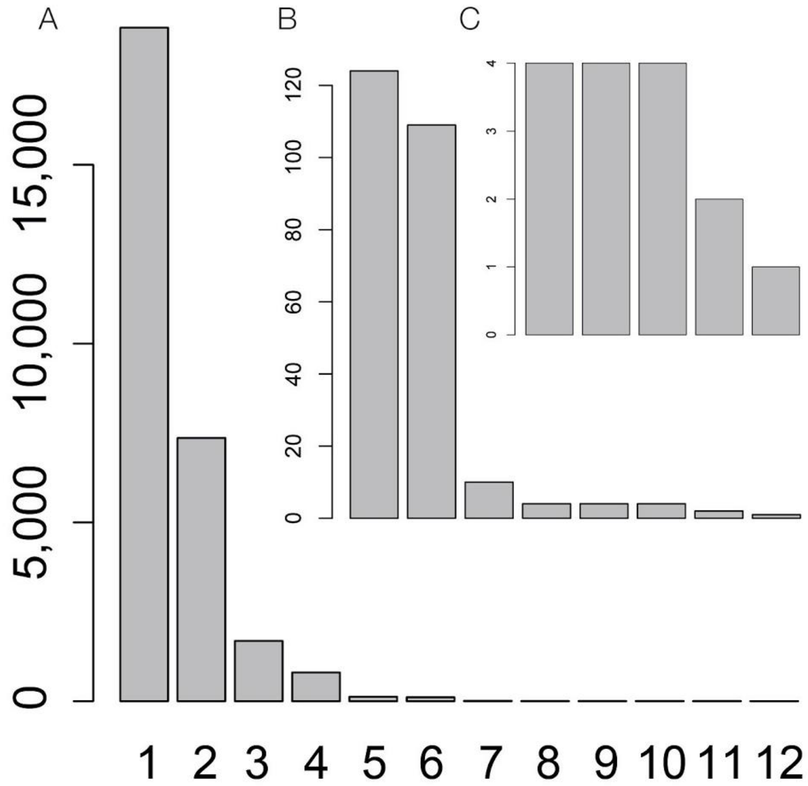


**Supplementary Figure 5.** Histogram showing the number of identified independent eQTL effects per QTL mapping round. A) Shows the number of eQTLs from round 1 to 12. Inlays B and C show the number of eQTLs identified in mapping round 5-12 and 8-12 respectively.

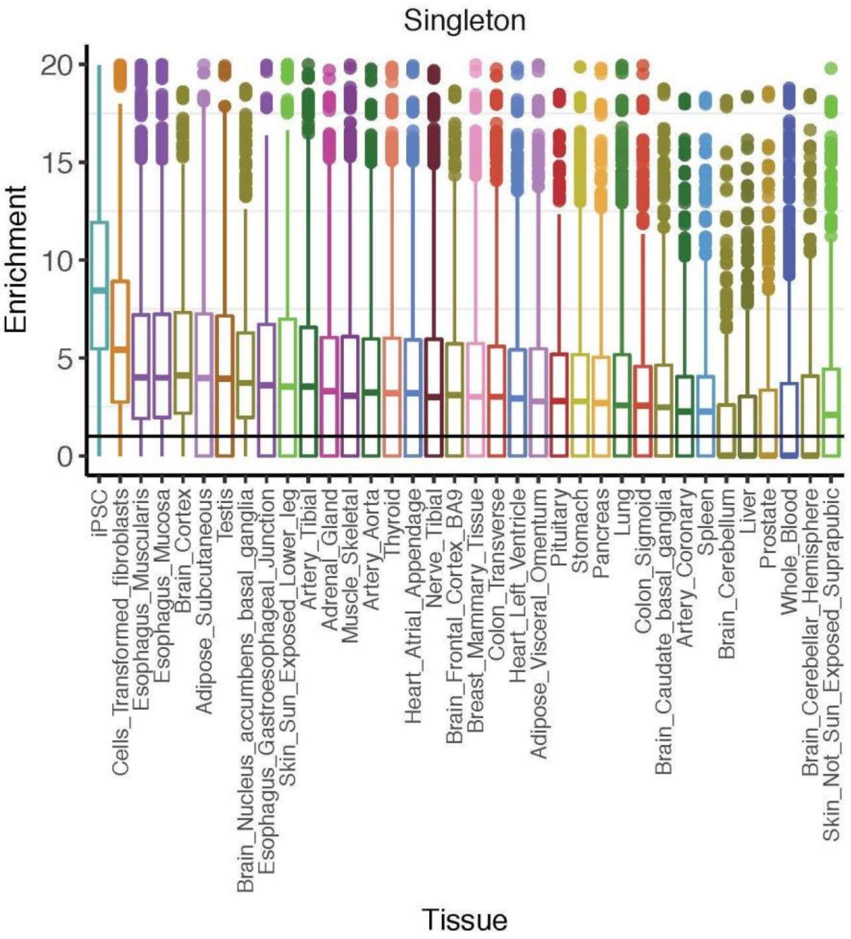


**Supplementary Figure 6.** Enrichment for rare, deleterious variants across iPSC and GTEx tissues across 10,000 permutations at fixed gene number

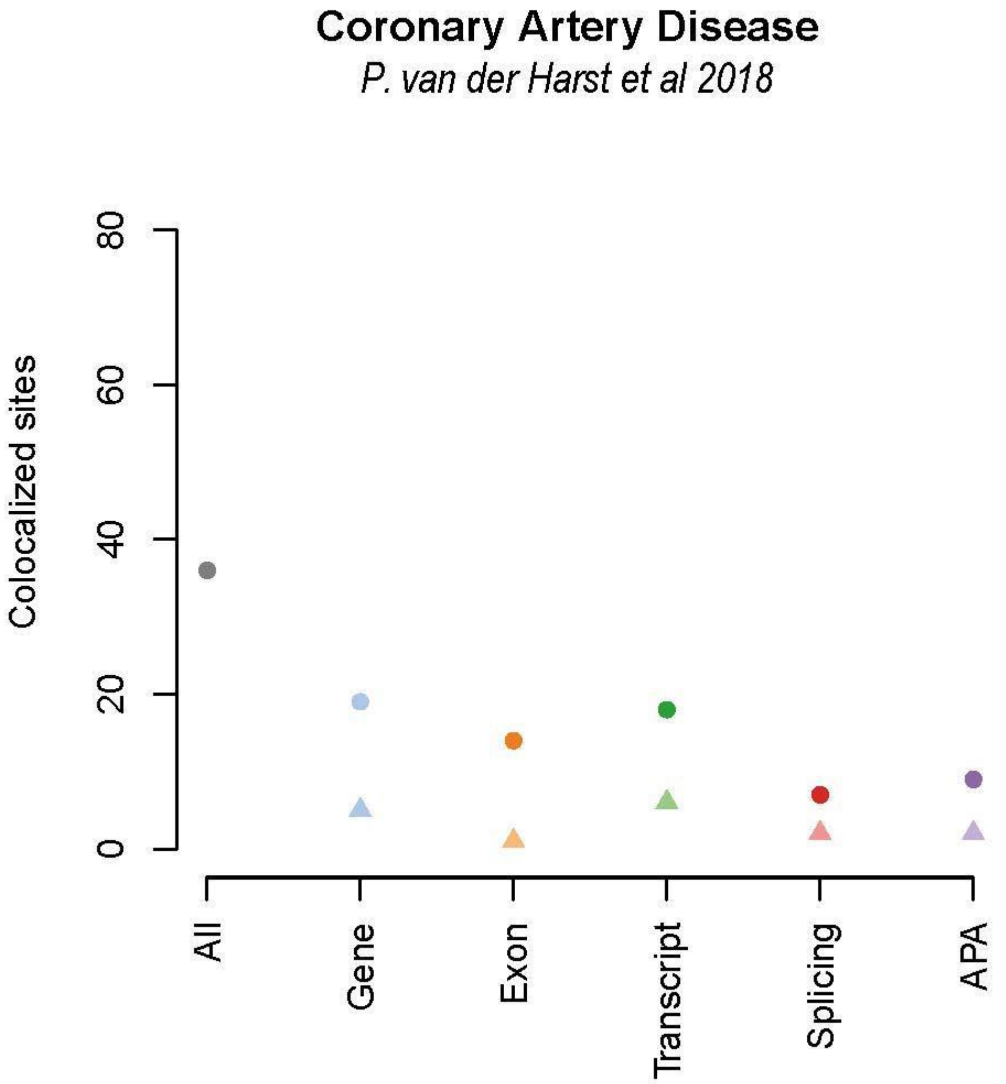


**Supplementary Figure 7.** Colocalization for the Coronary artery Disease GWAS by van der Harst et al 2018 and the *cis*-QTLs identified within i2QTL. Colocalizations between GWAS and QTL loci were found for in total 36 out of 93 loci (38.7%). Depicted are the total number of colocalizations (left in grey), and the colocalizations split out over the QTL types (the number of unique colocalized loci per type are marked with a triangle, and the round depicts all of the colocalized loci per QTL type).

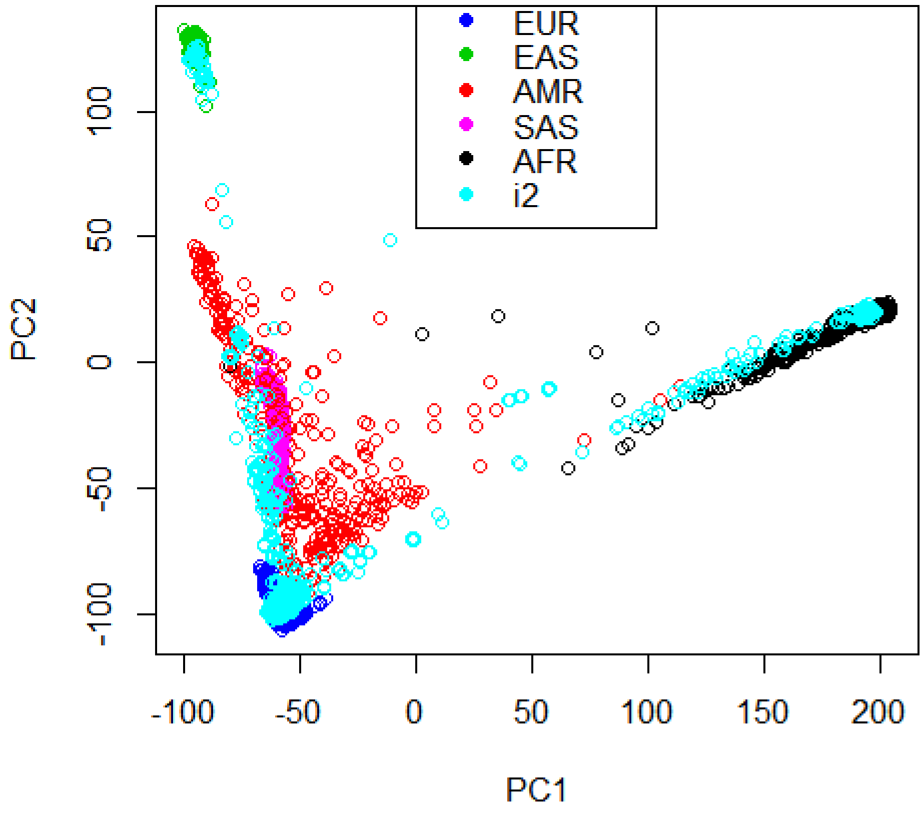


**Supplementary Figure 8.** Principal component analysis over the genotype data of the i2QTL studies projected into the 1000G data. Shown are the super populations as given by 1000G (Blue: European (EUR); Green: East Asian (EAS); Red: Ad Mixed American (AMR); Pink: South Asian (SAS); Black African (AFR); Cyan i2QTL samples).

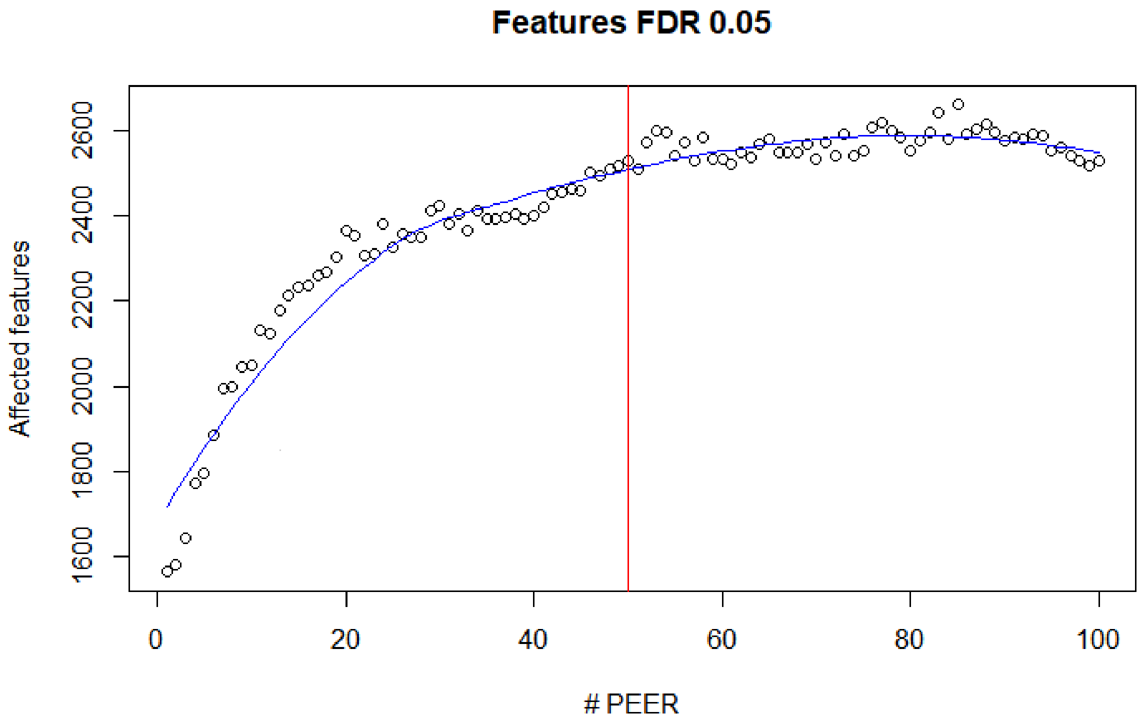


**Supplementary Figure 9.** The relation between the number of identified eQTLs vs the number of PEER factors corrected for. Shown are the selected number of PEER factors (50 in red) and the trend in terms of eQTL effects when regressing factors out of the expression data.

### Supplementary tables

Tables are available from:
https://drive.google.com/open?id=1zjdqCQVwX0IR8WZZDlSG_dvCHmlf8YqV

**Table S1. Overview of studies, data types, and study accession IDs for data used in the i2QTL study.** (Table_S1_SampleAccessionInformation.xlsx)
**Table S2. Metadata for i2QTL study samples**
(Table_S2_SampleMetadata.xlsx)
**Table S3. Gene-level cis-eQTL identified in the i2QTL study**
(Table_S3_GeneLevel_cisQTLs.xlsx)
**Table S4. Transcript-level cis-QTL identified in the i2QTL study**
(Table_S4_TranscriptLevel_cisQTLs.xlsx)
**Table S5. Exon-level cis-QTL identified in the i2QTL study**
(Table_S5_ExonLevel_cisQTLs.xlsx)
**Table S6. Splicing-level cis-QTL identified in the i2QTL study**
(Table_S6_SplicingLevel_cisQTL.xlsx)
**Table S7. APA-level cis-QTL identified in the i2QTL study**
(Table_S7_APA_Level_cisQTL.xlsx)
**Table S8. Outlier rare variant analysis across different outlier Z-score thresholds, gnomad MAF and CADD variant thresholds**
(Table_S8_OutlierRareVar.xlsx)
**Table S9. Rare variant enrichment for comparison analysis between i2QTL and GTEx v7**
(Table_S9_OutlierRareVarGTExComp.xlsx)
**Table S10. Variants associated with gene expression outliers in the comparative analysis of i2QTL and GTEx v7**
(Table_S10_OutlierIntersect.xlsx)
**Table S11. Top gene-level trans-eQTL identified in the i2QTL study**
(Table_S11_TopGeneLevel_transQTL.xlsx)
**Table S12. Gene-level trans-eQTL identified in the i2QTL study**
(Table_S12_GeneLevel_transQTL.xlsx)
**Table S13. Colocalizations identified using cis-eQTL from the i2QTL study**
(Table_S13_colocalizationResults.xlsx)
**Table S14. Colocalizations identified using trans-eQTL from the i2QTL study**
(Table_S14_trans_colocalizationResults.xlsx)
**Table S15. Overview of genes tested across different analyses in the i2QTL study**
(Table_S15_GeneTestingInformation.xlsx)

### Banner authors

Ordered by consortium, primary affiliation and last name.

#### HipSci


##### Wellcome Trust Sanger Institute

Chukwuma A. Agu, Alex Alderton, Shrada Amatya, Petr Danecek, Rachel Denton, Richard Durbin, Daniel J. Gaffney, Angela Goncalves, Reena Halai, Sarah Harper, Christopher M Kirton, Andrew Knights, Anja Kolb-Kokocinski, Andreas Leha, Shane A. McCarthy, Yasin Memari, Minal Patel

##### European Molecular Biology Laboratory

Ewan Birney, Francesco Paolo Casale, Laura Clarke, Peter W. Harrison, Helena Kilpinen, Davis J. McCarthy, Oliver Stegle, Ian Streeter

##### King’s College London

Davide Denovi, Ruta Meleckyte, Natalie Moens, Fiona M. Watt

##### University of Cambridge

Willem H. Ouwehand, Ludovic Vallier

##### University of Dundee

Angus I. Lamond, Dalila Bensaddek

##### University College London

Philip Beales

#### iPSCORE

##### University of California, San Diego

Angelo D. Arias, Paola Benaglio, Neil C. Chi, Matteo D’Antonio, Agnieszka D’Antonio-Chronowska, Christopher DeBoever, Margaret K.R. Donovan, Sylvia M. Evans, KathyJean Farnam, Kelly A. Frazer, Melvin Garcia, Lawrence S.B. Goldstein, William W. Greenwald, Olivier Harismendy, David A. Jakubosky, Kristen Jepsen, He Li, Hiroko Matsui, Thomas J. McGarry, Bradley C. Nelson, Daniel T. O’Connor, Fangwen Rao, Erin N. Smith, Gene W. Yeo

##### Salk Institute

W. Travis Berggren, Carl T. Dargitz, Kenneth E. Diffenderfer, Rachel Feiring, Juan Carlos Izpisua Belmonte, Veronica Modesto, Athanasia D. Panopoulos

#### GENESiPS

##### Stanford University

Ivan Carcamo-Orive, Joshua W. Knowles, Thomas Quertermous

#### PhiLiPS

##### University of Pennsylvania

Christopher D. Brown, YoSon Park
